# Structure of the TnsB transposase-DNA complex of type V-K CRISPR-associated transposon

**DOI:** 10.1101/2022.08.05.502904

**Authors:** Francisco Tenjo-Castaño, Nicholas Sofos, Blanca Lopéz-Méndez, Luisa S. Stutzke, Anders Fuglsang, Stefano Stella, Guillermo Montoya

## Abstract

CRISPR-associated transposons (CASTs) represent unique mobile genetic elements that co-opted CRISPR-Cas immune systems for RNA-guided DNA transposition. Type V-K CAST is composed by Cas12k, TniQ, TnsC and TnsB. Here, we present the 2.46 Å cryoelectron microscopy structure of the *Scytonema hofmannii* CAST TnsB transposase in complex with the strand transfer DNA in a post-catalytic state. The shTnsB strand transfer complex maintains the intertwined architecture of the MuA phage transpososome. However, the building of the assembly depends on different local interactions. The protein-DNA complex forms a pseudo-symmetrical assembly in which the 4 protomers of shTnsB adopt two different conformations. The recognition of the transposon ends is accomplished by two small helical domains. The two protomers involved in the strand transfer reaction display a catalytically competent active site composed by three acidic residues (DDE), while the other two, which play a key role in the complex architecture, show catalytic pockets where the DDE residues are not properly positioned for cleavage. Quantification of *in vivo* transposition assays of mutants in key DNA binding residues, reveals that the lack of specificity generally decreases activity, but it could increase transposition in some cases. Our structure sheds light on the strand transfer reaction of the DDE DNA transposases and offers new insights into RNA-guided transposition in CAST systems.

## Introduction

The discovery of an adaptive prokaryotic immune system called <u>C</u>lustered <u>R</u>egularly <u>I</u>nterspaced <u>S</u>hort <u>P</u>alindromic <u>R</u>epeats (CRISPR) in which the repeats associate with Cas (<u>C</u>RISPR <u>as</u>sociated) proteins, has constituted a revolution in life sciences. CRISPR-Cas systems are highly diverse ribonucleoprotein (RNP) complexes with different evolutionary origins. They are divided into two classes, Class 1 and Class 2, the former possessing a multisubunit effector complex and the later a single protein effector ^1^. The two classes are further divided into six types (I-VI) depending on the identity of the nuclease module, and many subtypes depending on which other Cas proteins are present in other functional modules. Especially, class 2 members have garnered much attention as they have been developed into versatile RNA-guided nucleases for RNA-guided genome editing, which has radically altered life sciences, enabling genome manipulation in living organisms ^2^.

Recently, several CRISPR-Cas machineries have been found associated with Tn7-like transposon systems in types I, IV, and V. These CAST systems^3,4 5^ are a product of an evolutionary process by which Tn*7*-like transposons seem to “parasite” the CRISPR-Cas system for transposon mobilization. These RNP complexes do not degrade their target DNA and operate exclusively in prokaryotes. They insert large DNA cargos (10–30 kb) at specific genome regions without the need for homology directed repair^4,6–8^, combining the site-selection precision of CRISPR-Cas with the integration properties of transposons^9^. Therefore, CASTs are thought to provide a very promising system for the development of nextgeneration gene-editing tools.

CAST I-F, I-B, and V-K subtypes, from *Vibrio cholerae* (vc), *Anabaena variabilis* (av) and *Scytonema hofmanni* (sh) respectively, were the first ones to be discovered^3,4^, but recent bioinformatic searches of metagenomic databases have vastly expanded the known CAST *repertoire* to over 1000 non-redundant subsystems representing Types I, IV, and V^7^. To date, all known CASTs are derived from Tn*7*-like transposons and include the corresponding crRNAs and *Cas* genes necessary for target selection^8, 39 7^, and the core transposition machinery within a Tn*7*-like transposon locus. This includes TnsB, TnsC, TniQ (a homologue of *E. coli* TnsD), and, in certain cases, TnsA genes. In an analogy with the Tn7 transposon systems, the CAST Tn*7* proteins are thought to assemble into a pre-integration nucleoprotein complex that involves TnsA (in Types I and IV), TnsB, TnsC and TniQ, to regulate transposition into the insertion site. TnsA is an endonuclease that cleaves the 5’-ends of the transposon^10^, and interacts with TnsB, TnsC and DNA^10–12^. TnsB is a recombinase and catalyses the cleavage of the 3’-ends of the transposon. In the canonical Tn7 system, the interaction of TnsA and TnsB is necessary to activate catalysis^13^. TnsC is part of the AAA+ ATPase family and directs TnsA/TnsB to the insertion site^12,14^. The interaction of TniQ with the target DNA bound by the CRISPR-Cas complex is thought to create a distortion in the DNA structure, allowing TnsC to recognize both TniQ and the DNA^15^, and subsequently leading to the insertion of the transposon at the attachment site. However, CAST type V-K systems lack TnsA in their loci, making the transposition mechanism unclear^16,17^, as it is unknown how the 5’-ends of the transposon are processed to proceed with the cargo integration.

TnsB transposases belong to the retroviral integrase superfamily with an RNase H fold catalytic domain and a DDE active site motif. TnsB associates with the left and right element ends of the transposon and catalyses their cleavage generating free hydroxyl groups, which are later used in a nucleophilic attack downstream of the target sequence^18^. Finally, it performs the strand-transfer reaction that leads to the insertion of the DNA cargo into the attachment site.

The shTnsB transposase shares homology with other members of the DDE integrase family such as *E. coli* TnsB, MuA and Tn5053 (Supplementary Fig. 1). The structure of *E. coli* (ec) TnsB in complex with the end of the transposon has provided new evidence explaining differences in the recognition of the left and right ends of the element ^19^. However, the sequence identity of ecTnsB with shTnsB is low, thus making it difficult to understand the details of the integration mechanism of Tn7/CASTs and other transposon systems containing integrases of the DDE family.

To comprehend how shTnsB facilitates RNA-guided transposition in the CAST V-K system, we determined the structure of the shTnsB-DNA complex captured after the strand transfer reaction by single particle cryo-EM at 2.46 Å resolution (Fig. 1). The structure reveals an entangled architecture of the shTnsB protein around the DNA building a pseudo-symmetrical assembly in which the 4 subunits of shTnsB can be grouped in two different conformations. The DNA in the DDE catalytic pockets displays a sharp bent after the strand transfer reaction. Site directed mutagenesis and *in vivo* transposition assays reveal the important role in transposition of key DNA-binding residues. The strand transfer reaction complex of shTnsB opens new avenues to understanding RNA-guided transposition in CAST systems.

**Figure 1.**
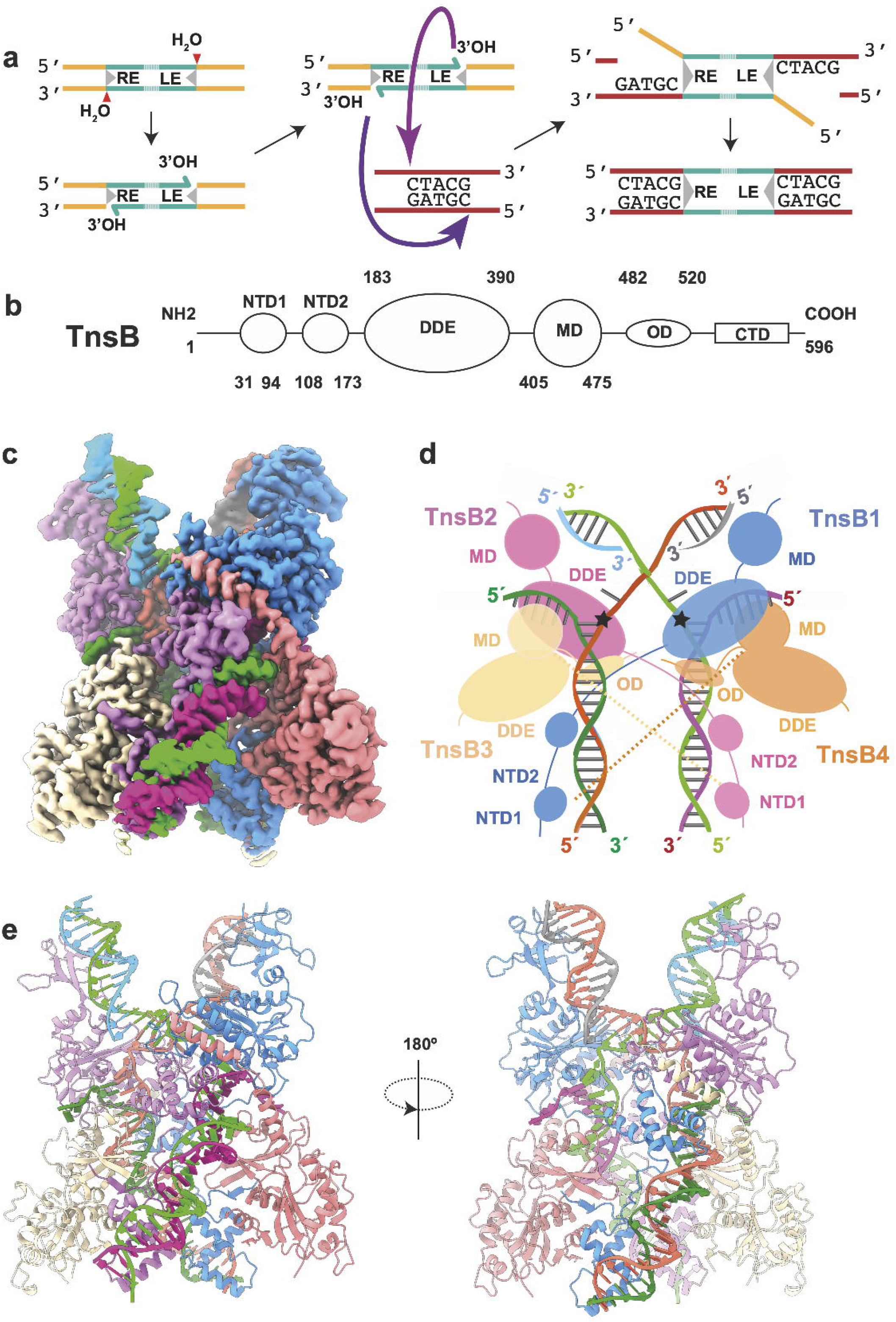
Cryo-EM structure of the CAST shTnsB subunit representing the post-catalytic state of the strand transfer complex. **a)** General scheme of the integration reaction followed by the TnsB family of transposases. At each end, the DDE active site catalyzes the attack of H_2_O at the target site (1). The direct attack of the 3’OH of the DNA cargo on the target DNA site promotes the strand transfer (2). Target binding leaves a 5 nucleotides spacer between the sites of attack. The 5’ overhangs are removed and filled in by a polymerase. The structure represents the strand transfer product (STC). **b)** Architecture of the shTnsB protein. The polypeptide is composed by an N-terminal domain that can be divided in two subdomains (NTD1 and NTD2), a catalytic DDE domain (DDE), a middle domain (MD), an oligomerisation domain (OD) and a C-terminal domain (CTD). **c)** Cryo-EM density map at 2.4 Å global resolution of the TnsB (STC) in a post-catalytic state (See also Supp. Figures 5-6, Supp. Table 2). The map is colored according to the cartoon describing the structure in panel d. **d)** Cartoon of the shTnsB STC. The complex is composed by 4 shTnsB protomers and 6 DNA oligonucleotides representing the post-catalytic state of the STC (Fig. 1a, stage 3). The dashed lines depict the interaction between the MDs and NTB1s due to the curvature of the DNA branches. **e)** Ribbon diagram displaying an overview of the shTnsB STC assembly.

## Results

### Isolation of shTnsB and generation of the strand transfer complex (STC)

The recombinant shTnsB protein was expressed in *E. coli* and purified using a combination of affinity and size exclusion chromatography (SEC). The protein behaved as a monomer in SEC-MALS displaying a MW of 68 kDa (Supplementary Fig. 2a-b, Methods). We analysed its DNA binding properties by EMSA using oligonucleotides with a different number of short and long repeats (SR and LTR respectively) present on the left and right elements (LE and RE) of the transposon end sequences (Supplementary Fig. 2c-d, Supplementary Table 1). The analysis of the band shifts with all the different substrates revealed a ladder of discrete bands, which increased with the concentration of the protein, suggesting the binding of one protein monomer per repeat. Assemblies higher than 6 proteins could not be observed as their size prohibited entry into the acrylamide gel (Supplementary Fig. 2c). Then, we tested whether shTnsB could bind RE or LE independently (Supplementary Fig. 2d). Similarly, the number of bands observed when shTnsB was mixed with RE corresponds to 5 TnsB binding sites. However, the number of bands observed when shTnsB was mixed with LE was 4, i.e., one higher than the expected 3 binding sites. This could be explained by the interaction of two shTnsB:LE complexes similar to the one observed in the STC complex (Fig. 1) or in other transposase structures not bound to the target DNA, such as in Tn5 ^20^. However, aggregation of the shTnsB:LE complex could also cause a band shift and cannot be completely discarded. Finally, we tested binding of shTnsB to dsDNA containing only LTR(6)-SR(1) (Supplementary Fig. 2e), and observed two bands corresponding to two shTnsB binding sites as expected. Furthermore, we incubated this complex in buffer containing different NaCl concentrations at 37 °C and 45 °C to determine whether these variables might change shTnsB affinity. Altogether, the EMSA results suggested that shTnsB binds every repeat present in both RE and LE in a cargo-independent way.

Initially, we prepared grids and collected data using a sample containing TnsB, and a RE and LE not bound to each other (Supplementary Fig. 2e). This sample was rather heterogeneous and suffered from preferential orientation. The processing of these data rendered low resolution reconstructions of insufficient quality to build an atomic model (Supplementary Fig. 3). However, two protuberances could be observed associated to an elongated density, suggesting the presence of two shTnsB protomers bound to a dsDNA face of a SR-LTR set. The elongated density assigned to the DNA is bent in a similar manner as in the recently published structure of the RE bound ecTnsB ^19^, further supporting that the low-resolution map corresponds to a pre-transposition complex.

To stabilize a shTnsB-DNA complex, we designed oligonucleotides to reconstitute the STC, i.e., the state representing post-transposition instead of pre-transposition as described above (Methods, Supplementary Table 1). The STC DNA consists of a LTR(8)-SR(5) and a SR(1)- LTR(6) dsDNA bound to the attachment site of CAST. A distance of 5 bp between the insertion sites of LTR(8)-SR(5) and SR(1)-LTR(6) was chosen as CAST transposition causes a 5 bp duplication in the insertion site ^21^ (Fig. 1a).

The reconstituted DNA STC was mixed with the purified protein and the assembly of the shTnsB-STC complex was verified by SEC-MALS (Supplementary Fig. 2b). Two peaks were observed, the lower MW peak contained the unbound protein and DNA used in the reconstitution, while the high MW peak, which eluted at 358.7 kDa, coincided very well with the association of the STC DNA (104.6 kDa) and 4 protomers of shTnsB (68 kDa), whose theoretical MW is 376.6 kDa. This assembly was subjected to single particle analysis by cryo-EM, which rendered a 2.46 Å resolution map where we built an atomic model of the shTnsB-STC complex in its post-catalytic state (Fig. 1a)

### Overall structure of shTnsB-STC complex

The shTnsB protein is composed of 6 different domains (Fig. 1b) and resembles the architecture of Mu transposase (MuA) ^22^, even though sequence identity is limited to the catalytic DDE domain (Supplementary Fig. 1a). We collected 4728 movies and set coordinates for 9.6 million particles, which were reduced to 415k after 2D classification. Using this set of particles we performed non-uniform refinement, 3D variability analysis and subsequently heterogeneous refinement in cryoSPARC ^23^. This approach yielded two maps at 2.5 and 2.8 Å. Further rounds of refinement and 3D variability analysis using the 260k particles of the 2.5 Å map rendered a cryo-EM map at 2.46 Å global resolution that allowed the modelling of the shTnsB-STC complex (Fig. 1c, Supplementary Fig. 4-5, Supplementary Table 2 and Methods). However, the large flexibility of the DNA compromised the visualisation of the DNA ends containing SR(5) and SR(1), preventing the determination of possible contacts with shTnsB (Supplementary Fig.4-5, Supplementary Video 1 and Supplementary Table 1).

The structure of the shTnsB-STC complex resembles an elongated X with curved arms. The longer transposon end DNAs forming the inferior longer branches and the bent target DNA the superior shorter ones (Fig. 1d-e, 2a). The four protomers of shTnsB (shTnsB1-4) display an intertwined assembly on the STC DNA. The protein is found in two different conformations to associate with the STC DNA and build the complex (Fig. 2b). shTnsB1 and 2 depict an extended conformation along the branched nucleic acid structure, where we could not observe the OD and CTD domains (Fig.1b, d). In both protomers each of the NTD1 and NTD2 domains are associated with the inferior end of the LTR(8) and LTR(6) elements respectively, while the catalytic DDE domain is visualized in the junction between the target DNA and the transposon ends of the opposite branch (Fig. 1c, 2b). On the target DNA side, the assembly is stabilized by interactions of the backbone with the MD and DDE domains of the shTnsB1 and 2 protomers (R416, Q427, K290, N428) (Fig. 2b). Thus, shTnsB1 accomplishes DNA recognition in the LTR8 end and its DDE domain catalyses the attack of the 3’OH in the LTR(6) branch on the target DNA, promoting the strand transfer reaction. A symmetric arrangement of the STC DNA is observed for shTnsB2, which catalyses the strand transfer of the LTR(8) on the other strand of the target DNA (Fig. 1a, Fig. 2b).

**Figure 2.**
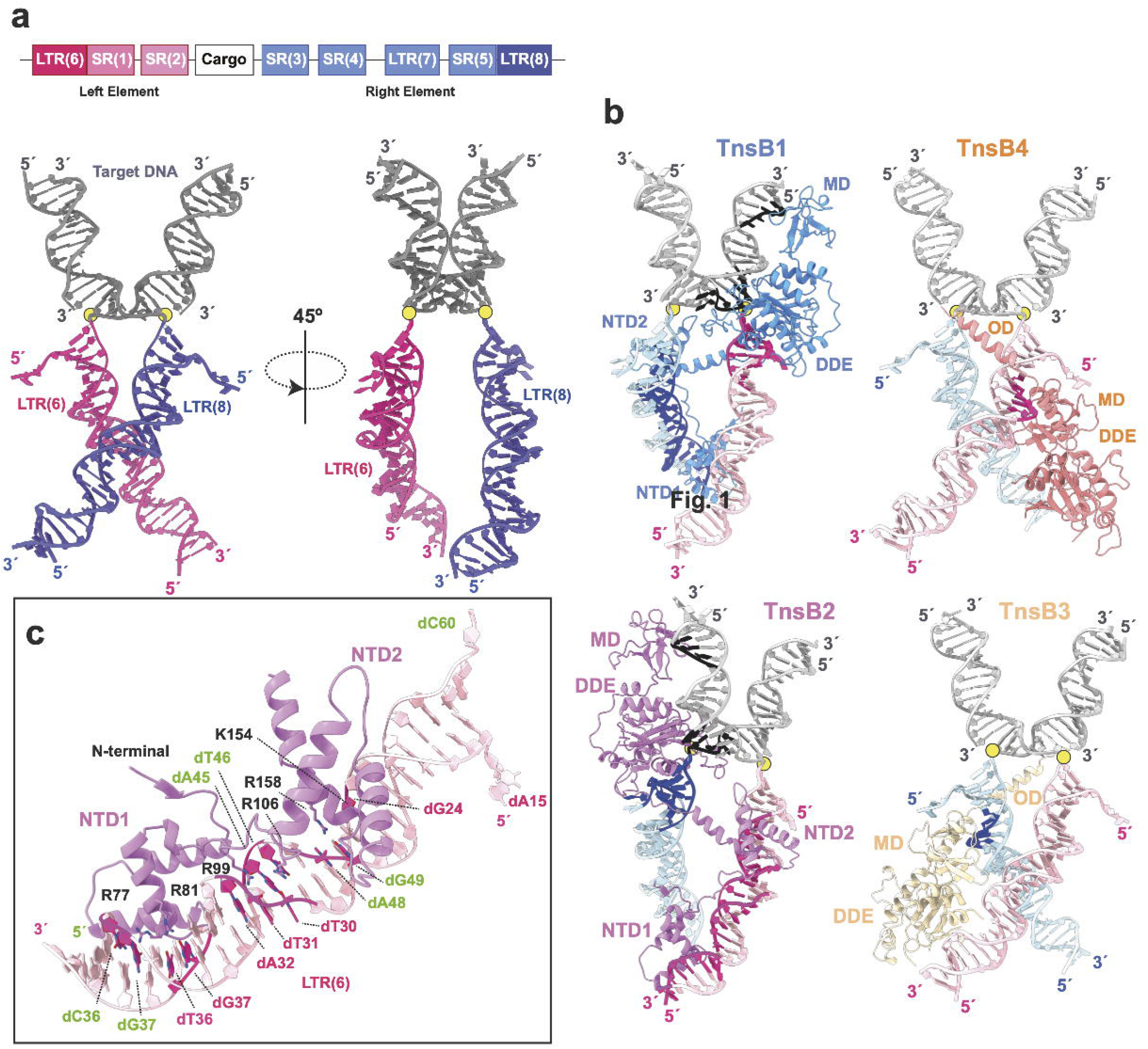
Architecture of the shTnsB STC complex. **a)** The upper panel shows a scheme of the *S. hofmanii* transposable element with the left and right elements (LE and RE) at each side of the DNA cargo. The strong color tone shows the region of each element present in the structure. The lower panel omit the protein moieties in the structure and shows the STC DNA. The DNA is colored according to the upper scheme for the LE and RE, while the region of the target DNA is shown in grey. The yellow dots indicate the site where the direct attack of the 3’-OH on target DNA has occurred to promote the strand transfer reaction. **b)** Detailed view of the shTnsB protomer association with the STC DNA. Two different associations are observed for the shTnsB molecules. The different sections of the DNA interacting with the proteins are shown in the corresponding strong colour tone. **c)** Detailed view of the helical NTD1 and 2 domains showing the specific interactions with the DNA bases in the LTR(6) region. The labelling of the nucleotides follows the colour code in Fig 1d-e (see also Fig. 1d and Supp Fig. 7).

The shTnsB3 and 4 protomers are arranged in a different conformation with few contacts with the backbone of the DNA (Fig. 2b). The NTD1 and NTD2 domains of these protomers are not observed in our structure, suggesting that there are highly mobile. In addition, the DDE domain is pulled out of the DNA while the MD accommodates the flap regions from the LTR6 and LTR8 ends after catalysis. The structure revealed the important role of these two protomers in the association of shTnsB1 and shTnsB2 with the DNA, as in this conformation they show an ordered OD domain. This helix is not observed in shTnsB1-2, but in the conformation adopted by shTnsB3-4, the helix facilitates the assembly of the STC complex by intertwining the NTD2 of shTnsB1 and shTnsB2, with the DDE domain of the opposite protomer in each branch of the DNA, thereby, linking the recognition of the two transposon ends with the catalytic sites of shTnsB1-2 (Fig. 1d, 2b).

### shTnsB DNA recognition

The recognition of the LTR(6) and LTR(8) is performed by the NTD 1 and 2 helical domains of the shTnsB1 and 2 subunits (Fig. 2c). The NTD1 domain structure reveals a helix-turn-helix Myb/homeodomain-like fold. Direct base contacts by the major groove binding helix principally account for the sequence-specific recognition. The well-conserved R77 makes polar contacts with dG37, while R81 associates with the bases of the consecutive dG36 and dT37 nucleotides on the complementary strand of the LTR6 in the LE (Supplementary Fig. 6, 7 and Supp. Table 1). Other residues interacting with the DNA (R58, R66, K84 andT78) show polar interactions with the backbone of both DNA strands. The NTD1 is joined to NTD2 by a long loop that runs along the groove of the DNA. The NTD2 domain is functionally similar to the second DNA binding domain in other transposases. However, it displays a limited conservation, and its structure is not related to any of the closest homologues of shTnsB (Supplementary Fig. 1). The well-conserved R99 in that loop builds polar interactions with the bases of dT31 and dA32 and dA45, while R106 also recognizes the bases of the dT30 and dT46 in different stands. The rest of the associations of the loop with the DNA involve backbone contacts. The NTD2 domain also binds to the DNA in the major groove. However, it displays fewer specific contacts with the bases. Only R158 and K154 associate with dA48-dG49 and dG24 respectively, while the rest of the residues associate with the backbone of the nucleic acid. These interactions are similar in the NTD1 and 2 domains on the LTR(6) and LTR(8) branches of the protein-DNA complex (Supplementary Fig. 6-7).

### Assembly of the protein-DNA complex

The NTD2 domains of the shTnsB1-2 subunits are connected to the DDE domains by a long loop. The intertwined conformation of these protomers with the DNA allows shTnsB1, which recognizes the DNA in LTR(8), to catalyse the strand transfer reaction on the LTR(6) end and vice versa for shTnsB2 (Fig 1c, 2b). This crossed positioning of the DDE catalytic domains is favoured by the conformation of the shTnsB3-4 molecules, which stabilize the assembly by intercalating a long bipartite helix that assembles the NTD2 and the DDE domains of different subunits. The adjacent MD accommodates the 5’flaps generated after the strand transfer reaction (Fig. 1d, Fig. 3a-c). The C-terminal helix of the OD is snugly fitted between the DDE domain of shTnsB2 and the NTD2 of shTnsB1 by a network of contacts combining polar and non-polar interactions (Fig. 3b), being the former at the end (E360-K520) middle (R367-D512-Q508) and final (S505-H272) regions of the helix. Another strong polar interaction is observed between the side chains D514 in the OD and R137 in the NTD2. A short loop joins this section with the OD N-terminal helix. The interaction of the second helix of the OD with the DDE domain is stabilized by non-polar interactions, while a group of residues in the MD of shTnsB3 (R416, Q425, N428) together with R174 and R179 in shTnsB2, form an electropositively charged region stabilizing the bases of the 5’overhang (Fig. 3c). Finally, the MD and DDE domains of the shTnsB3 protomer associate with the NTD1 of shTnsB2, mainly by a group of non-polar interactions. Interestingly, the N-terminal β-strand of shTnsB2 is inserted as an additional strand in the antiparallel β-sheet of the MD (Fig. 3d). Additional side chain polar interactions (E422-K169, N462-K102, and E54-H283) complete this association. Overall, the associations described here are very similar in the case of shTnsB4 with shTnsB1 and 2. Interestingly, we note that no interaction is observed between shTnsB3 and 4 in the shTnsB-STC complex.

**Figure 3.**
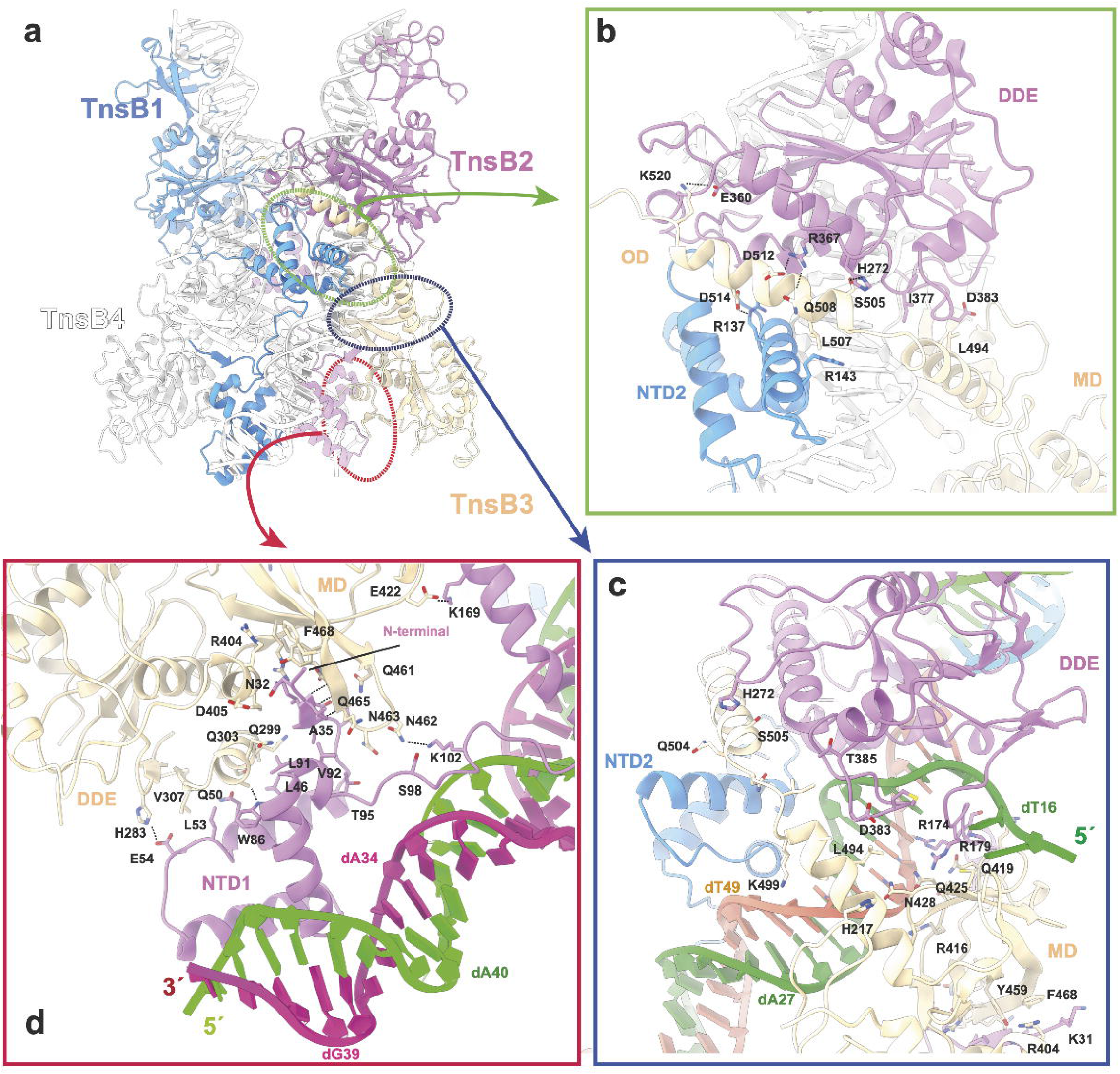
Assembly of the shTnsB protomers. **a)** Overview of the architecture of the shTnsB protomers and the interactions between the two types of conformations observed for the protein on one of the branches of the STC DNA (transparent). The colored ovals indicate the zoomed regions in the other panels. **b-c)** Detailed view of the interactions of the OD of shTnsB3 with the NTD2 and DDE domains of shTnsB1 and 2 respectively. **d)** Zoom of the LTR(6) region depicting the association of the N-terminal β-sheet of NTD1 in the MD of shTnsB3.

Collectively this network of interactions along the X shaped complex suggest that the different conformation of the shTnsB3 and 4 protomers promote an architecture of the assembly which favours a crossed interaction, thus associating the recognition of the LTR(6) and LTR(8) DNA sequences by the shTnsB1 and TnsB2 protomers with the DDE catalysis in the opposite branch.

### The DDE catalytic domain

The transposases from several superfamilies possess a catalytic domain containing an acidic amino acid triad (DDE or DDD). The DDE catalytic domain has been identified in the transposases from 11 of the 19 currently recognized superfamilies ^24^. This protein module catalyses the transposition reaction, whereby the element is excised from the donor site and inserted in the genome or in a mobile genetic element. Mutations in these catalytic residues have shown their critical role in transposition in the DDE domains of Tn5 and Tn10 ^25,26^ transposases. Integration of the elements into a new genomic location usually generates a short target-site duplication from host sequences (2–10 bp). In shTnsB the DDE motif consists of two aspartic acid residues (D205, D287) and a glutamic acid (E321) residue, located in a conserved core that forms a characteristic RNase H-like fold combining α-helices and β-strands (Fig. 4a, Supplementary Fig. 7). The 3D structure of the DDE triad forms a catalytic pocket which associates with divalent metal ions that assist in the various nucleophilic reactions during DNA cleavage. The DDE domains of the 4 shTnsB molecules in the assembly superimpose very well (rmsd 0.59 Å rmsd over 160 Cα). However, the DDE pocket displays two different arrangements depending on the conformation of the shTnsB protomer in the STC complex (Fig. 1). The DDE pockets of shTnsB3 and 4 do not contact the DNA and the catalytic residues of these subunits and are not properly positioned for the cleavage reaction (Fig. 4b). In the case of shTnsB1 and 2, they display a DDE catalytic pocket after the strand transfer reaction has been accomplished. An extra density in the pocket can be visualized (Fig. 4a), which could be assigned to H_2_O or Mg^2+^. E321 is 6 Å away, as the structure is captured in the post-catalytic state. The distances of the extra density to D205, D287 and the backbone phosphates suggest that the density could be assigned to the H_2_O molecule generated after the 3’-OH attack of one of the strands of the transposable element to the target DNA (Fig. 1a, 4a, Supplementary Figure 7).

**Figure 4.**
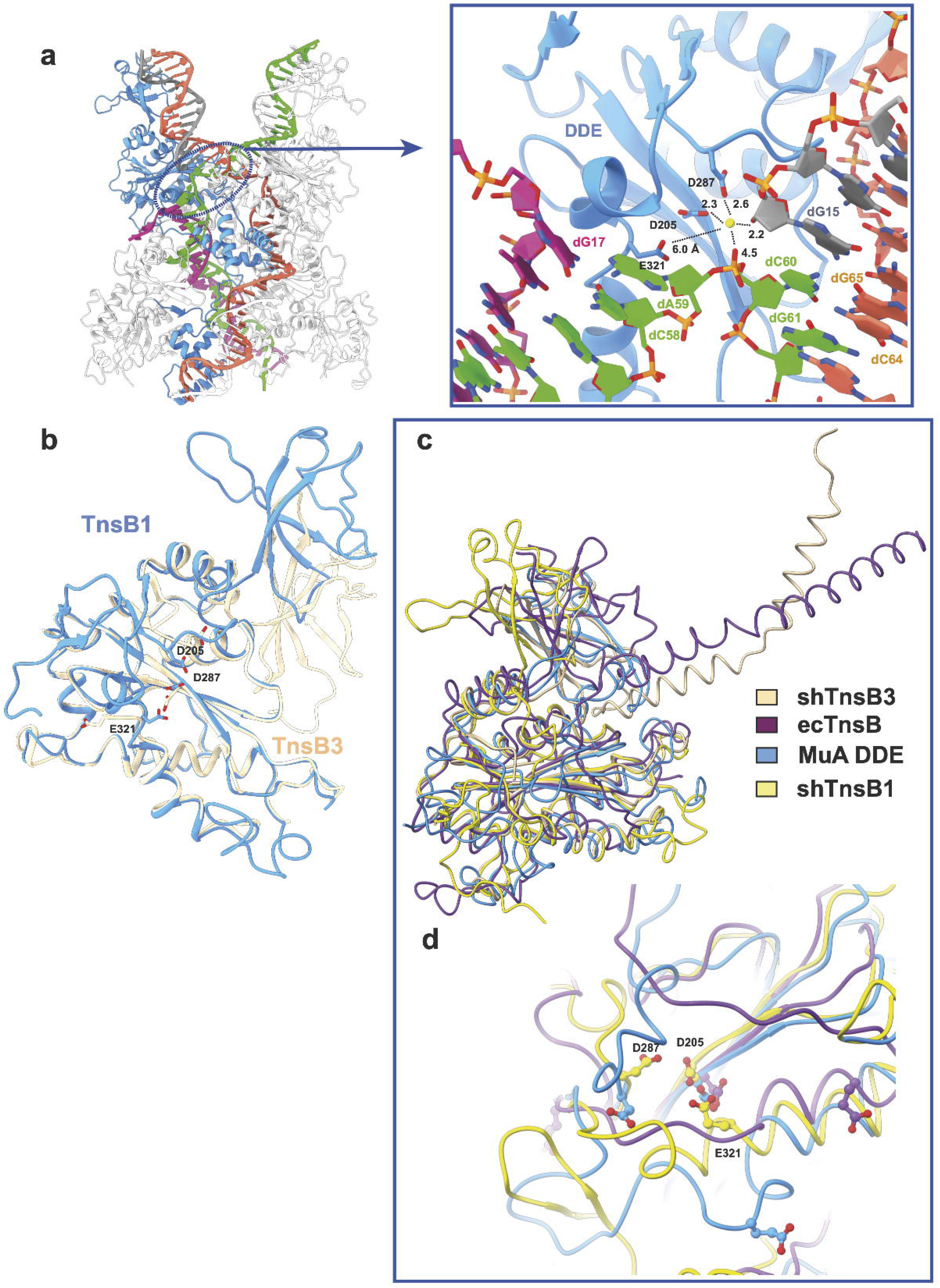
The DDE catalytic site of shTnsB. **a)** The encircled region locates one of the catalytic centers in the shTnsB-STC complex. The zoom of the region shows the conformation of the catalytic site after catalysis. The putative water molecule and the distances to the neighboring catalytic residues and the DNA are shown in Å. **b)** Superposition of the DDE domains of the two conformations found in the protein DNA complex for shTnsB. **c)** Superposition of the DDE domains of shTnsB2 and 3 with ecTnsB (PDB:7pik) and the MuA core domain (PDB:1bco). The RMSDs for the superposition are in the range between 0.7 to 1.2 Å for 160 to 250 Cα. **d)** The lower panel shows that despite the high structural similarity of the DDE domains, the only subunit in all these structures that displays a competent catalytic pocket is shTnsB2, which is the only one associated with DNA.

Despite sequence differences, the recent structure of the Tn7 transposon ecTnsB transposase ^19^ end recognition complex provides a close functional homologue for comparisons. The DDE domain is the best conserved module of shTnsB, and a search for shTnsB DDE homologues using DALI ^27^ revealed the domains of ecTnsB and the phage Mu transposase as the closest homologues, together with different viral integrases (SRV, HIV-1, HTLV-1). The comparison of the two shTnsB conformations with these proteins showed that the core of the domain superimposed very well with other integrases (Fig. 4c), but only the catalytic pocket of shTnsB2 showed the acidic amino acid triad at the appropriate distances to catalyse the strand transfer reaction. In the rest of the cases, the catalytic residues were not properly positioned (Fig. 4d). Noteworthy, the ecTnsB protomers in the end recognition complex displayed a conformation similar to shTnsB3 and 4 protomers in the STC. However, in the latter, the DNA binding domains of these two subunits were not visualized (Fig. 1-2), suggesting that the high flexibility observed in the distal sections of the STC DNA (Supplementary Video 1) disturb their binding to the SR(1) and SR(5) regions of the LE and RE.

Collectively, the structural comparison suggest that the DDE domain maintains a very well conserved 3D structure, however, the comparison of different structural homologues suggest that the architecture of the catalytic pocket of the transposase is properly arranged once it associates with the target DNA.

### DNA binding by shTnsB, ecTnsB and MuA transposases

The shTnsB protein display homology to ecTnsB and MuA transposases. However, besides the DDE domain few residues are conserved between them (Supplementary Fig. 1). These differences are especially evident in the DNA-binding domains where multiple sequence alignments from several canonical Tn7 TnsB proteins and diverse classes of CAST elements have shown substantial differences ^19^. The NTD1 of shTnsB shows significant 3D similarities with the HTH domain of the DBD1 in ecTnsB (RMSD 2.4 Å for 66 Cα) and the Iβ domain of MuA (RMSD 2.8 Å for 62 Cα). Nonetheless, the presence of the N-terminal β-strand (N32-T36) is exclusive of shTnsB, and this region plays an important role in building the assembly of the STC complex (Fig. 3d). This structural feature is not observed in the MuA STC complex ^22^, the only structure of a prokaryotic transposase STC complex available so far (Fig. 5).

**Figure 5.**
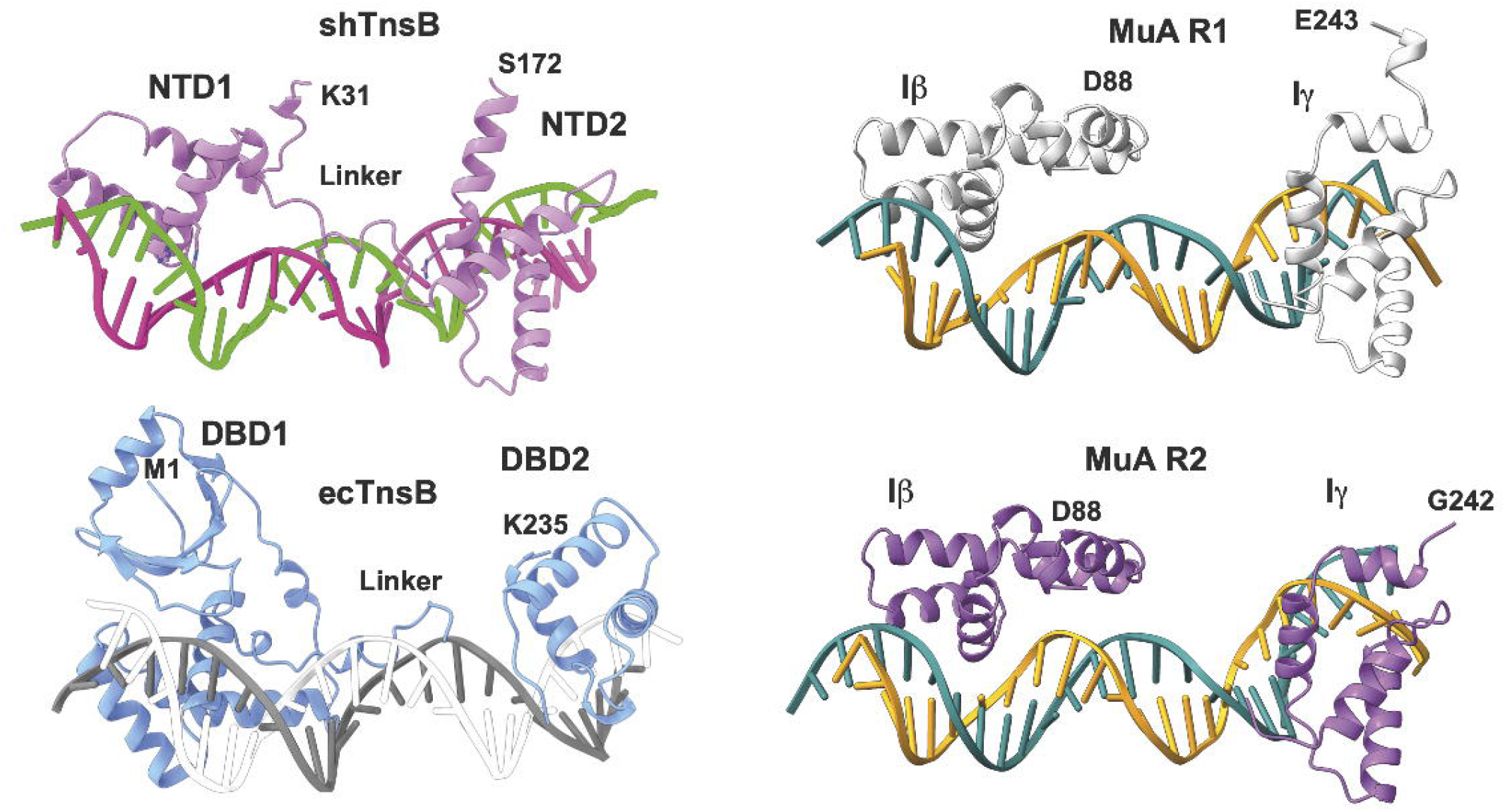
Comparison of the DNA binding domains of shTnsB, ecTnsB and MuA. The DNA binding domains of the three different transposases are shown aligned on the DNA (PDB:8AA5, 7PIK and 4FCY respectively). In the case of MuA the same domains bound to the different repeats are shown. For shTnsA only the domains of the protomers 1 and 2 were built. In the case of ecTnsB the DNA binding domain of one of the subunits in the end complex is depicted.

Interestingly, the NTD2 domain of shTnsB does not show 3D similarity to its counterpart domains in ecTnsB and MuA transposases (Fig. 5). Instead, NTD2 superimposes well with Pax6, a transcription factor containing a bipartite paired DNA-binding domain (RMSD 2.8 Å for 60 Cα), which has critical roles in development of the eye, nose, pancreas, and central nervous system ^28^. The extended linker joining NTD1 and 2 is partially visualized in ecTnsB and it is not observed in the MuA DNA binding domains (Fig 5). This linker makes minor groove contacts, and the carboxy-terminal helix-turn-helix unit makes base contacts in the major groove, as observed in the shTnsB-STC complex (Fig. 5).

Together, the comparison suggests that the TnsB proteins share a common overall DNA-binding mode, including two domains that recognize the different SR and LR elements in the assembly. However, the domains involved in binding undergo different structural variations to accomplish the specific protein-DNA interactions for each transposase. This diversity could be required to accommodate the sequences of a large number of Tn7-like elements, preventing transposases from recognizing DNA sequences belonging to other transposons^29^.

### The STC complexes

The ecTnsB end-complex structure has provided information on the recognition of the transposon ends ^19^, but so far, there is no structural information on their STC complex. However, the conformation of the ecTnsB subunits in the end complex resemble the arrangement adopted by the DDE domain of the shTnsB3 and 4 protomers in the shTnsB-STC complex. However, the MD and OD domains positions are oriented differently (Fig. 4c). Nonetheless, this conformation is rather different than the architecture of the protomers 1 and 2 of shTnsB, which are involved in the catalysis of the strand transfer reaction (Fig 2b). The conformation of these subunits is more similar to its counterparts in the MuA-STC complex (Fig. 6a).

**Figure 6.**
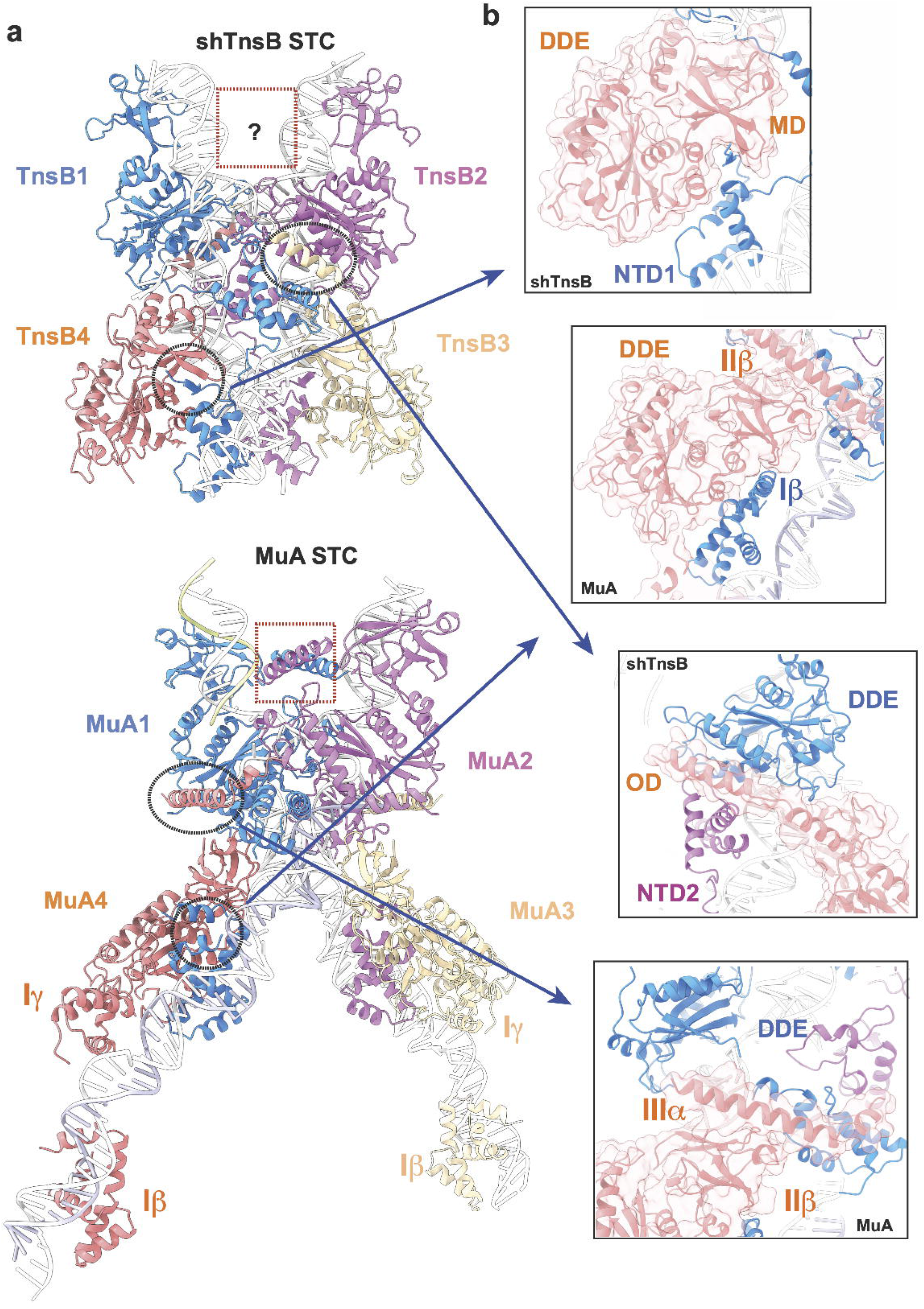
Analysis of the Strand Transfer complexes of shTnsB and MuA. **a)** The shTnsB-STC and MuA-STC complexes are shown following the same color scheme as in Fig. 1, except for the DNA which is colored in white. **b)** The encircled regions on both complexes are compared to show key differences in the assembly of the STC complexes.

Although, the sequence of shTnsB is distantly related to the Mu phage transposase MuA (Supplementary Fig. 1), the general association on the MuA-STC complex shares similitudes with the CAST protein (Fig. 6a). However, the assembly also presents important differences. One of the main differences is that in the MuA-STC complex, the DNA binding domains of the subunits involved in the assembly (MuA 3 and 4) are bound to the corresponding repeats in the LE and RE (Fig. 6a), while in the shTnsB-STC they could not be modelled, suggesting that they are highly flexible. This notion is supported by the high variability observed in the distal part of the X shaped DNA structure (Supplementary Video 1). The fact that those regions of the MuA-STC structure ^22^ are involved in crystal contacts could reduce the flexibility, thus facilitating the binding of the Iβ and Iγ domains to the repeats in the LE and RE ends.

Interestingly, the association of the unique NTD1 N-terminal β-strand in shTnsB is not observed in MuA (Fig. 6b), as this secondary structure element is not present neither in the Iβ domain of MuA nor in the DBD1 of ecTnsB. This observation suggests that in the ecTnsB-STC complex this association cannot occur. Another important difference arises from the assembly of the complex, the OD domain in shTnsB intercalates, with multiple interactions, between the NTD2 and the DDE catalytic domain building the association between shTnsB proteins. In the case of MuA the association between the 4 protomers in that region of the STC complex is different, displaying considerably less interactions between the different domains (Fig. 6b). Furthermore, the OD domains are not observed in the protomers 1 and 2 of the shTnsB-STC complex, while in the case of MuA the helices in the IIIα domain are crossed in the interior of the V shaped target DNA region (Fig. 6a). The possibility that the interaction between these helices in the MuA structure is due to crystal contacts could not be disregarded.

### The effect of DNA binding in transposition

To validate the shTnsB:DNA interactions in the STC, we tested the transposition activity of a series of substitution mutants where conserved amino acids displaying polar interactions with the DNA were replaced by alanine (Fig. 7, Supplementary Fig. 1). We mutated R77, R81 (both present in NTD1), R99 (present in the linker between NTD1 and NTD2), R158 (present in NTD2), R188, R223, K290 and R380 (present in the DDE domain) (Fig. 7b). The R77A, R81A and R158A mutants showed a dramatic effect on on-target transposition activity, suggesting that shTnsB can no longer recognize the repeats in RE and LE (Fig. 7b). Interestingly, the nucleotides involved in this interaction (Supplementary Fig. 6) are conserved between the different repeats further supporting importance of this interaction in donor DNA recognition ^21^. As for R99, even though it is present in a disordered region of shTnsB, it has an intimate interaction with the phosphate backbone and the bases of the LTRs. These nucleotides are also conserved within the RE/LE repeat set ^21^. However, the effect of its substitution was variable with replicates displaying either higher or lower activity than the wild type. In the case of R223A and R380A, the activity was also affected. These residues seem to be involved in stabilizing the post-transposition state. R223 interacts with the central nucleotides of the 5 bp sequence between the nicks generated by transposition, while R380 interacts with the non-cleaved transposon ssDNA positioned out of the complex. Finally, despite its conservation and proximity to the attachment site, mutants R188A and K290A did not abolish on-target transposition. Our results confirm the functional importance of these residues involved in LTR recognition and STC stabilization and reveal possible sites for improvement of the transposase properties for genome engineering purposes.

**Figure 7.**
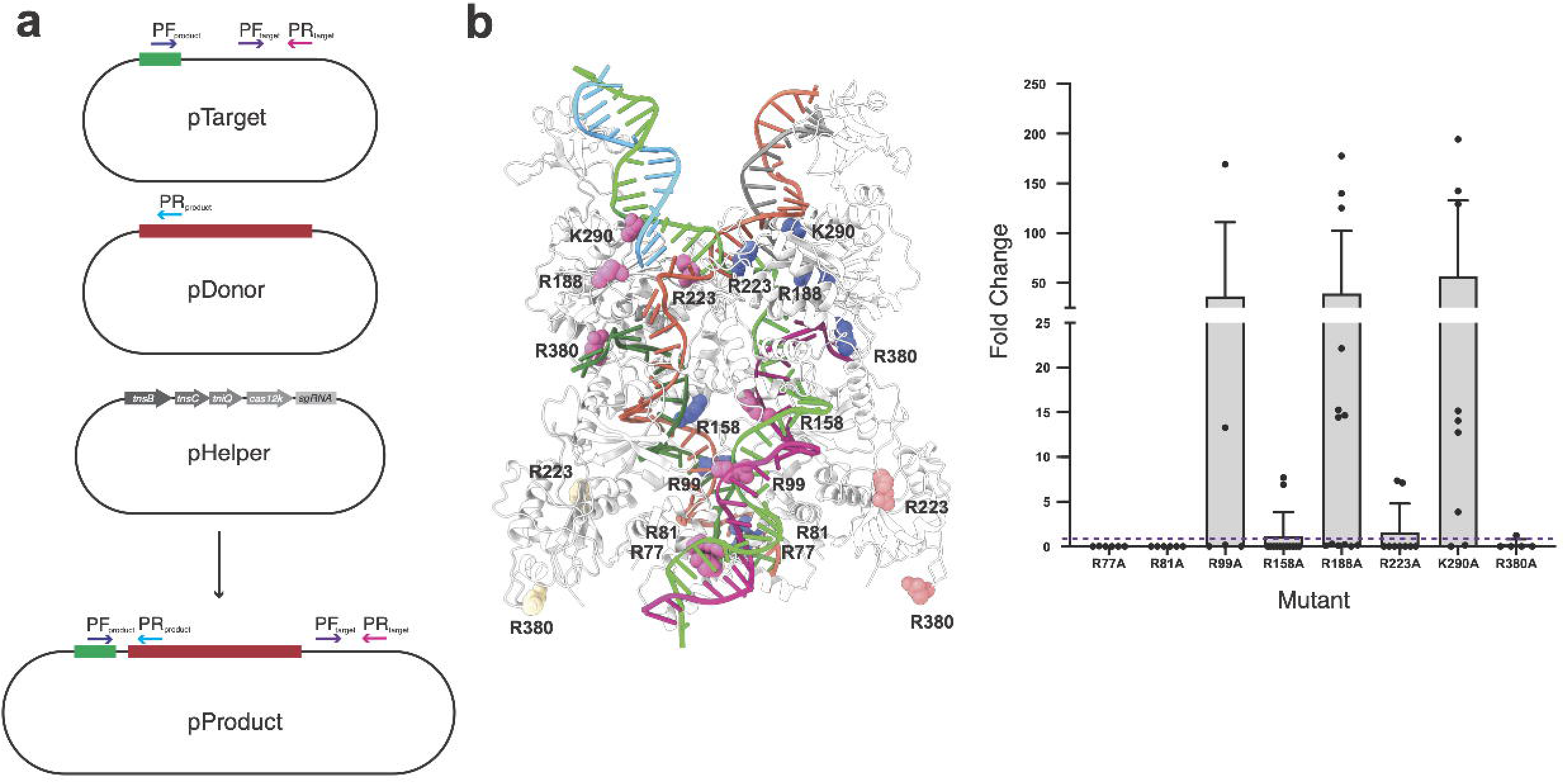
*In vivo* transposition assay and qPCR quantification. **a)** Transposition was carried out by transforming bacteria with a plasmid containing a target site, shown in green, recognized by Cas12k:sgRNA (pTarget), a plasmid containing a donor DNA with the CAST RE and LE sequences shown in red (pDonor) and a plasmid comprising the sgRNA, Cas12k, TnsC, TniQ and TnsB coding sequences (pHelper). Integration of the donor sequence into pTarget was quantified by qPCR using a set of primers amplifying a region of pTarget where no transposition occurs (PF_target_ and PR_target_) and a set of primers that amplify a region between the target and the donor LE sequences (PF_product_ and PR_product_). **b)** Position of selected amino acids within the STC to be mutated and tested for transposition activity. **c)** Activity fold-change measured by qPCR for each TnsB mutant. Bars represent the average and whiskers represent the standard deviation. Individual data points are also depicted. The wild type activity is represented by the dashed line.

## Discussion

Recent studies have described new CRISPR-Cas systems that associate with Tn*7* like-transposon proteins^17–19 7^. These CAST systems recruit the CRISPR-Cas machinery and “parasitize” on it to direct DNA integration at a specific locus^4^. CASTs combine the site-selection precision of CRISPR-Cas with the ability to integrate large DNA cargos via the Tn*7* machinery. Prokaryotic CAST systems lack the adaptation system (Cas1-Cas2) and Type I Cas3 nuclease required for target DNA degradation. It is not known how their crRNA is matured, but processed crRNA assembles an RNP, enabling crRNA-guided DNA binding but incapable of interference. The Type V-K CAST is a minimal system composed of a Cas12 related protein that builds an RNP with a crRNA and three Tn7-related proteins: TniQ, TnsC and TnsB. The type V-K system of *S. hofmannii* catalyses RNA-guided DNA transposition by unidirectionally inserting segments of DNA downstream of the protospacer, and it can integrate DNA into targeted sites in bacterial genomes^21^. Therefore, CASTs expand our understanding of the functional diversity of CRISPR-Cas systems and establishes a paradigm for precision DNA insertion. The inactive CRISPR-Cas effector complexes in the major part of the CAST systems belong to the Class 1 CRISPR systems, which assemble into big multisubunit complexes. The main exception so far is Type V-K, which represent one of the most promising systems for the development of gene editing tools due to the reduced size of the inactive effector Cas12k, composed by a single protein (639 residues) and a sgRNA (254 nt).

Last year witnessed a large increase of structural knowledge of CASTs systems ^30–33^. Structural information of TnsC, TniQ and Cas12k of type V-K^34–36^, as well as for type I-F Cascade complex bound to TniQ^37,38^ is available. However, despite these advances, there are key components involved in transposition and their complexes where we have no detailed molecular information. Furthermore, we need precise insight of the sequence of interactions and intermediates leading to precise DNA cargo insertion to address a possible engineering of the CAST systems. This lack of thorough understanding of how transposition is accomplished by the different CASTs, precludes their repurposing for precise insertion of DNAs into gene editing tools.

In this manuscript we have determined the structure of *S. hofmannii* TnsB protein of Type V-K CAST. The only component, where no structural information was available so far. The low resolution shTnsB-end complex (Supplementary Fig. 3) suggests that the transposon end recognition is very similar to the ecTnsB ^19^; however, the high heterogeneity and flexibility of the assembly did not allow a detailed molecular characterisation of the DNA recognition. This agrees with experiments in the Tn7 system which have shown that the pre-transposition complex is less stable than the post-transposition complex and that transposon protection changes between the pre- and post-catalytic stages ^39,40^. In addition, EMSA experiments suggest that shTnsB can interact with the RE and LE in the pre-catalytic state in the absence of the target DNA (Supplementary Fig. 2). Thus, shTnsB interaction with the transposon ends seems to be independent on the presence of other components from the CAST system such as TnsC, which was previously shown to interact with TnsB ^34^.

The higher stability of the post-catalytic complexes led us to assemble the shTnsB-STC complex, which we were able to determine at high-resolution. This complex represents the stage after completion of the transesterification reaction with the target DNA, revealing its 3D assembly in the post-catalytic state. The overall structure resembles the architecture of the phage MuA transposome ^22^. Yet, key differences between the assemblies can be observed especially in the NTD2 domain (Fig. 6). One possible similitude between the shTnsB and MuA, which could differentiate them from the Tn7 system, is that the Type V-K and Mu could use replicative transposition, which requires the cleavage of only one DNA strand, while Tn7 performs a cut-and-paste reaction, involving the cleavage of the non-transferred strand by TnsA. Therefore, the structural similarities of the overall STC complex between shTnsB and MuA would be supported by these mechanistic similitudes. Yet, we will expect that the Tn7-STC complex should accommodate TnsA. From our structure it is not clear how the 5’-ends could be processed in the Tn7 system.

The post-catalytic shTnsB-STC structure revealed very few base-specific protein-DNA contacts (Supplementary Fig. 6), as it has been also observed in the ecTnsB pre-catalytic complex ^19^. The NTD1 domain of shTnsB partially shares its architecture with its counterparts in ecTnsB and MuA, while NTD2 is unique in the sense that it shares structural homology with the Pax6 transcription factor (Fig. 5). A singular structural feature of shTnsB is the N-terminal strand which is used to link the recognition in one of the transposon arms with the catalytic centre on the opposite site and vice versa, whether this feature exerts some allosteric regulation in the DDE catalytic site remains to be determined.

While there is structural information on the DDE catalysis from other transposase families (Ref Tn5 structure etc etc), the shTnsB structure sheds light on the DDE catalytic pocket of the TnsB family, as the resolution of the MuA transpososme hindered a detailed study of the catalytic site ^22^. The two subunits of shTnsB engaged in the reaction display the acidic triad in a post-catalytic state. The attack of the 3’-OH of the transposon strands with the 5’of the target DNA leaves uncoupled bases (dT62 and dA63) in each strand and a water molecule bound to the aspartic residues and the DNA backbone (Supplementary Fig. 6). Based on the structures of the shTnsB-STC and the ecTnsB end-complex structures, we can conclude that the active pocket of the TnsB proteins is not arranged for catalysis before the target DNA is bound (Fig. 4d). Most likely this is a mechanism to preclude non-specific cleavage.

Tn7, and CAST types I-B and I-F rely on TnsA and TnsB to carry out transposon integration^49,41^ As previously mentioned, TnsA is not present in CAST Type V-K systems. Therefore, the formation of TnsA-TnsB-TnsC assembly, which occurs in the T7 elements, and is supposed to occur also in other CASTs types, should not be present in *S. hofmannii* Type V-K. Nevertheless, the C-terminus of shTnsB (Fig. 1b) contains and unstructured segment that interacts with shTnsC ^34^, as in the case of the T7 system ^42^, suggesting that at least this two proteins form an assembly indicating where the integration is occurring. However, TnsA’s absence in type V-K leads to further questions regarding integration intermediate resolution since non-cleaved transposon 5’ ends result in cointegrate transposition products^43,44^. Transposons relying on DDE transposases have different mechanisms to solve this problem^45–47^. For instance, Tn5053 contains resolvase TniR which uses *res* sites, also present in the transposon, to resolve the cointegrate^48,49^ while other transposases generate hairpins to excise the remaining DNA^50^. Additionally, DNA replication can solve the cointegrate in the case of replicative transposition^47,50^. How type V-K CAST deals with cointegrate resolution is still unknown. However, certain facts suggest it might need a host-specific resolvase. First, even though type V-K CAST is present as a single copy in its natural host, when transposition is carried out in *E. coli*, 20% of insertion products are cointegrates, pointing away from the possibility of TnsB creating hairpins at the ends of the transposon DNA^44^. Second, the V-K system gene set (TnsB, TnsC and TniQ) is similar to that of Tn5053’s, which contains TniA, TniB (TnsB and TnsC homologs) and TniQ^44^. It could be hypothesised that a resolvase not encoded in the V-K transposon locus but still present in the natural host could have the same activity as resolvase TniR. Further studies would need to be performed in *S. hofmanni* (or other related cyanobacteria) before we can fully understand RNA-guided transposition on Type V-K. To solve these questions is a key premise to successfully generate new engineered CAST complexes able to integrate large DNA cargoes in eukaryotic genomes.

Our structure provides a mechanistic understanding of how CAST shTnsB recognizes the DNA sequences in the RE and LE in the left and right ends of the transposon in the STC. The structure also shows the conformational gymnastics of the shTnsB protein to assemble with the DNA and provides a detailed view of the active pocket after catalysis. Full elucidation of the assembly mechanisms of shTnsB with the rest of the CAST components will be required to harness CAST systems for RNA-guided genome engineering.

## Acknowledgements

We thank the Danish Cryo-EM National Facility in CFIM at the University of Copenhagen and specially Tillmann Pape for support during cryo-EM data collection. We also thank the Protein Expression Unit at CPR for assistance in protein expression and purification. FJTC. is a member of the Copenhagen Bioscience PhD programme. G.M. is part of the Novo Nordisk Foundation Center for Protein Research (CPR), which is supported financially by the Novo Nordisk Foundation (grant NNF14CC0001). This work was also supported by grant NNF0024386, grant NNF17SA0030214 and Distinguished Investigator (NNF18OC0055061) grants to G.M, who is a member of the Integrative Structural Biology Cluster (ISBUC) at the University of Copenhagen.

## Author contributions

GM conceived the study, FTC and GM designed the biochemical experiments. FTC and SS set up the expression and purification protocol. FTC and AF characterized the protein-DNA binding properties. FTC, and LS created the mutants and performed transposition experiments. FTC and BLM performed the SEC-MALS analysis, FTC prepared EM grids and collected the cryo-EM images with NS. FTC and NS performed cryo-EM data processing, built the models and GM proceeded with cryo-EM map and structure analysis with their help. The global results were discussed and evaluated with all authors. GM coordinated and supervised the project and wrote the manuscript with input from all the authors.

## Methods

### Protein production and purification

The *Scytonema hofmannii* (UTEX 2349) *tnsB* coding sequence (IDT) was cloned with a *C-* terminal hexahistidine (His)-tag into pET-21 vector (Supplementary Table). Proteins were expressed in *E. coli* BL21 pRARE cells. *E. coli* cultures were grown at 37 °C in Terrific Broth (TB) medium with 100 mg/L ampicillin to an OD_600_ of 0.6. Overexpression of proteins was induced with 250 mM of IPTG for 16-18 h at 18 °C. Cells were harvested by centrifugation, flash frozen in liquid Nitrogen, and stored at −80 °C. Pellets were subsequently resuspended in lysis buffer (50 mM HEPES pH 7.5, 500 mM NaCl, 5 mM MgCl_2_, 1 mM TCEP, 1 tablet of Complete Inhibitor cocktail EDTA Free (Roche) per 50 mL, 50 U/mL Benzonase, 2 mg per 50 mL lysozyme). The resuspended pellets were stirred for one hour at 4 °C. Lysis was completed by sonication and lysate was centrifuged at 10000 X g for 45 min to separate the soluble and insoluble fractions. The soluble fraction was loaded into a 5 mL HisTrap FF Crude column (Cytiva) equilibrated in buffer IMAC-A (50 mM HEPES pH 7.5, 500 mM NaCl, 1 mM TCEP, 10 mM imidazole, 1 tablet of Complete Inhibitor cocktail EDTA Free (Roche) per 500 mL), and bound proteins were eluted by stepwise increase of the imidazole concentration with buffer IMAC-B (50 mM HEPES pH 7.5, 500 mM NaCl, 1mM TCEP, 500 mM imidazole, 1 tablet of Complete Inhibitor cocktail EDTA Free (Roche) per 500 mL). Enriched protein fractions were pooled together and NaCl concentration was reduced to 200 mM using dilution buffer (50 mM HEPES pH 7.5, 50 mM NaCl, 1 mM TCEP). The sample was applied onto a HiTrap Heparin HP column (GE Healthcare) equilibrated with buffer Heparin-A (50 mM HEPES pH 7.5, 200 mM NaCl, 1 mM TCEP). The protein was eluted with a linear gradient of 0–100% buffer Heparin-B (50 mM HEPES pH 7.5, 1 M NaCl, 1 mM TCEP). Fractions containing the protein were pooled and mixed with TEV-protease to cleave the his-tag. Then, the protein was concentrated and further purified by size exclusion chromatography (SEC) using a HiLoad 16/600 Superdex 200 column (Cytiva) equilibrated in SEC buffer (50 mM HEPES pH 7.5, 500 mM NaCl, 1 mM TCEP). Fractions containing pure protein were pooled, concentrated to 9 mg/mL, flash-frozen in liquid nitrogen and stored at −80 °C.

### Electromobility Shift Assay (EMSA)

DNA (donor, RE, LE or LTR(6)-SR(1)) was mixed with TnsB in buffer containing 50 mM HEPES pH 7.5, 300□mM NaCl, 5 mM MgCl_2_ and 1□mM TCEP. DNA and TnsB concentrations are indicated in Supplementary Figure 3c-e. Samples were incubated at 37°C for 30 min and loaded onto 6% DNA retardation gels (Thermo Fisher Scientific). DNA was detected by staining the gel with GelRed (VWR). The same gel was afterwards stained with InstantBlue (Life Technologies) to detect the presence of TnsB and compare band shifts.

### DNA production

DNA sequences corresponding to individual repeats, double repeats, and STC ssDNA were purchased from IDT. DNA molecules containing three or more repeats were obtained by PCR with a donor DNA cloned into pUC57 (Genewiz) as template (Supplementary Table 1). Complete donor DNA was amplified using primers 5’-tgctacgtctctacgtgtacagtgactaat-3’ and 5’ ccacaaaaggcgtagtgtacagtgacaaat-3’, RE was amplified using primers 5’-tgctacgtctctacgtgtacagtgactaat-3’ and 5’-atatacaaagcgacagtcaatttgtcattatgaaaat-3’, and LE was amplified using primers 5’-attatattgatgacatttaatttgtcatcaattaattaag-3’ and 5’-ccacaaaaggcgtagtgtacagtgacaaat-3’. The amplified sequences were purified using QIAquick PCR Purification kit (Qiagen) and eluted in nuclease-free water.

### Cryo-EM Sample Preparation

The precatalytic complex was prepared using 1 μM of RE, 1 μM of LE and 8 μM of shTnsB in 300 μL of buffer containing 50□mM HEPES pH 7.5, 300□mM NaCl, 5 mM MgCl_2_ and 1□mM TCEP. The sample was incubated for 30 minutes at 37°C and concentrated to 50 μL in 10 kDa filters. The STC was reconstituted as follows. The LE and RE ssDNA were mixed separately, incubated at 95°C for 5 minutes, and left on ice for 10 min. Subsequently, the corresponding target ssDNA was added to RE and LE. Both samples were incubated separately at 70°C for 5 minutes and left on ice for 10 minutes. Finally, the RE and LE were mixed and incubated at room temperature for 10 minutes. Finally, 50□mM HEPES pH 7.5, 300□mM NaCl, 5 mM MgCl_2_ and 1□mM TCEP were added to 10 μM DNA STC DNA before adding 40 μM TnsB. The sample was incubated for 30 minutes at 37°C.

Grids for both samples (precatalytic complex and STC) were prepared by applying 3 μL of freshly prepared STC (1:2 dilution) to UltrAuFoil 300 mesh R1.2/1.3 holey grids (Quantifoil), glow-discharged for 60 s at 10 mA (Leica EM ACE200), and plunge-frozen in liquid ethane (pre-cooled with liquid nitrogen) using a Vitrobot Mark IV (FEI, Thermo Fisher Scientific - blotting time of 3 s, 100% humidity, 4 °C).

### Cryo-EM data processing

Data processing was carried out using cryoSPARC version 3.3.2 ^23^. Patch motion correction and patch CTF estimation was performed. 9,646 K particles were initially picked from 4,574 micrographs using the blob picker with min and max diameters of 50 and 200 Å respectively, and subsequently extracted, and sorted by two rounds of 2D classification. The selected 2D classes contained 415 K particles, extracted using a 416×416 pixel box. An *ab initio* 3D reconstruction was used as a starting volume for non-uniform refinement, and the resulting particles were analyzed by 3D variability analysis ^51^ (Supplementary Figure 4), which revealed a continuous conformational variability in the distal ends of the DNA and nearby shTnsB domains (N-terminal part and CTD of chains A and B). Two of the most spatially different volumes produced were used as references for heterogeneous refinement and were further refined by non-uniform (NU) refinement to a global resolution of 2.47 Å (269 K particles) and 2.81 Å (155 K particles), based on the gold-standard Fourier shell correlation (GSFSC) 0.143 cutoff criterion (Supplementary Figure 5). Per-particle defocus and per-group CTF parameters were refined during the NU-refinement. On visual inspection, the density of the 2.47 Å map appeared considerably more continuous. This map was further improved in the distal ends by another iteration of 3D variability analysis of the particles ^51^ and using the most spatially different volumes as references for another round of heterogeneous refinement. Most particles ended up in one class (208 K) which were finally refined by NU refinement to a 2.46 Å map. 3DFSC analysis revealed a sphericity of 0.928 and an angular resolution range of 2.4-3.1 Å (Supplementary Figure 5). Local resolution analysis was performed with MonoRes ^52^ using the cryoSPARC wrapper, which revealed a range of ~2-10 Å with the majority being around 2.5 Å, and distal part being 4-8 Å. Model building was performed in maps that had been post-processed using DeepEMhancer ^53^. Half-maps were used as input, while normalization and masking/denoising of the input volumes was performed automatically in the tightTarget mode, yielding maps of significantly improved interpretability.

### Atomic model building and refinement

An initial model was generated based on the MuA structure (PDB:4fcy), using the automated protein structure homology-modelling server, SWISS-MODEL ^54^. It was rigid body fitted into the map using ChimeraX ^55^. The model fit was inspected and further fitted in COOT ^56^, and less ordered N- and C-terminal regions were removed. An Alphafold ^57^ model of the TnsB monomer was subsequently used to fit some of these regions into the map as well. The model was further rebuilt *de novo* in COOT, and refined using phenix.real_space_refine ^58^, while the Namdinator molecular dynamics flexible fitting server was used to speed-up a nd improve the model to map fit and for model refinement in both early and late stages of refinement ^59^.

#### *In vivo* transposition assay and mutagenesis

shTnsB mutants (Fig. 7) were generated in the pHelper vector using the In-Fusion cloning kit from Takara and primers containing the desired point mutations. CAST transposition activity was detected using vectors pHelper, pTarget and pDonor obtained from Addgene (Cat# 127922, 127926 and 127924). shTnsB mutants (Fig. 7) were generated in the pHelper vector using the In-Fusion cloning kit from Takara and primers containing the desired point mutations. In vivo assays were performed following Saito *et al.* ^60^. Briefly, 12 ng each of pHelper, pTarget, and pDonor were transformed into 30 μL of TransforMax EC100 pir+ Electrocompetent *E. coli* (Lucigen). Cells were recovered for 1 hour and plated on LB media containing ampicillin, kanamycin, and chloramphenicol. Cells were scraped 24 hours after plating, lysed in 15 μL colony lysis buffer (TE with 0.1% Triton X-100), boiled for 5 min, diluted with 30 μL of water and spun at 4000 g for 10 min to pellet debris. Supernatant was used for subsequent analysis.

#### Activity assay qPCR

The frequency of insertions relative to the concentration of the target plasmid was determined by qPCR (TaqMan Fast Advanced Master Mix, Applied Biosytems) with a set of primers/probe (5’-6-FAM, Int ZEN, 3’-Iowa Black, IDT) corresponding to an unmodified region of pTarget (primer forward: 5’-cgacagcatcgccagtcactatg-3’, primer reverse: 5’-caagtagcgaagcgagcaggac-3’, probe: 5’-tgcgttgatgcaatttctatgcgcacccgt-3’) and a set of primers/probe (5’-6-FAM, Int ZEN, 3’-Iowa Black, IDT) corresponding to the transposition product (primer forward: 5’-ggttgagaagtcatttaataaggccactgttaaacg-3’, primer reverse: 5’-aacgctgatgggtcacgacg-3’, probe: 5’-ctgtcgtcggtgacagattaatgtcattgtgac-3’) (Fig. 7). Each reaction (11 μL) contained 2 μL of extracted nucleic acids, 900 nM forward primer, 900 nM reverse primer, and 250 nM of probe. Fluorescence was measured using the LightCycler 480 Instrument (Roche). Data was analyzed by the 2^-ΔΔCt^ method normalizing with pTarget Ct.

## Supplementary Materials

### Supplementary Figure Legends

**Supplementary Figure 1.- Sequence alignment of representative members of the DDE transposase family**. The amino acid sequences of EcTnsB, shTnsB, MuA and Tn5053 were aligned by Clustal Omega (http://www.ebi.ac.uk/Tools/msa/clustalo). The figure was prepared with ESPript (http://espript.ibcp.fr). Residue numbers are labelled according to shTnsB sequence. Similar residues are shown in red and identical residues in white over red background. The different structural domains of shTnsB and their amino acid composition are depicted as boxes above the sequences and labelled with the same names and colour code as in Fig. 1a. The mutations performed in key residues commented through the text are marked with an asterisk.

**Supplementary Figure 2.- Purification and characterisation of shTnsB and the Strand Transfer Complex. a)** SDS-PAGE showing the purified shTnsB **b)** SEC-MALS of purified TnsB (green), STC DNA (orange), and TnsB:STC (purple). The results agree with the theoretical molecular weight of a TnsB monomer (68 kDa) and the separate target bound RE and LE DNA (52.6 kDa and 52 kDa). As for the complex, the observed molecular weight (358.7 kDa) corresponds to a complex containing the target bound RE and LE and a TnsB tetramer, which has a theoretical molecular weight of 376.6 kDa. **c)** Complexes containing donor DNA with two different cargos. Donor DNA concentration was 100 nM in all samples. TnsB concentration in each sample is indicated in the figure. **d)** Complexes containing only RE or LE. Donor DNA concentration was 300 nM in all samples. TnsB concentration in each sample is indicated in the figure. **e)** Complex containing only LTR(6) and SR(1) tested at different temperatures and NaCl concentrations. DNA concentration was 2 μM and TnsB concentration was 4 μM.

**Supplementary Figure 3.- Low resolution reconstruction of the shTnsB bound to RE/LE in the precatalytic state. a)** Representative micrograph of this protein-DNA complex. **b)** 2D classes displaying a bent DNA molecule bound to two protomers of shTnsB. **c)** 3D reconstruction based on selected 2D classes. Two protomers of shTnsB were fitted into the density based on the STC structure and following a similar pattern to the precatalytic structure of ecTnsB ^19^. NTD1 and NTD2 are shown in dashed circles indicating where the LTR would be.

**Supplementary Figure 4.- Single Particle Cryo-EM Analysis of the TnsB-STC. (a)** Representative cryo-EM micrographs of the TnsB-STC in vitreous ice on an UltrAuFoil 1.2/1.3 grid at −1.7 μm defocus. **(b)** Reference-free 2D class averages selected for *ab inito* reconstruction. **(c)** Overview of the cryo-EM data processing workflow as performed in cryoSPARC.

**Supplementary Figure 5.- Resolution Assessment and Validation of the TnsB-STC Cryo-EM map (a)** Local resolution maps in two orientation related by a 90° rotation around the y-axis. The inset shows the high resolution obtained in the central cut through the map, the clipping plane indicated above. **(b)** Gold standard Fourier shell correlation (GSFSC) curve after FSC-ask auto-tightening. **(c)** Orientation distribution plot (top) and posterior position directional distribution plot (bottom). **(d)** Histogram and directional FSC plot. Histogram (blue) shows the percentage of per angle FSC at the given spatial frequency. Global FSC (red) and ± 1 S.D. from mean directional FSC (green). Sphericity equals 0.928 out of 1. **(e)** Angular resolution map, and corresponding directional FSC volume (grey), in three perpendicular orientations. The relative angular resolution is indicated by the color bar.

**Supplementary Figure 6.- Protein-nucleic acids interactions in the shTnsB-STC complex.** Polar and non-polar contacts of the nucleic acids with the protein side and main chain are indicated (see figure key).

**Supplementary Figure 7.- Fit of ShTnsB-STC atomic model to the cryo-EM map.** Map regions are displayed at a 2 Å range within the displayed protein residues and nucleotides, at contour level 0.04 for protein-only panels, and level 0.1 for protein/DNA panels. Key residues and nucleotides are labelled. **(a)** OD α2, residues 504-520. **(b)** OD α1, residues 484-499. **(c)** Stacking interaction between W178, from the NTD2-DDM loop region, and dT16, from the overhand of the LE_polyA strand. **(d)** β-strands interaction, residues 32-36 and 464-468 of two neighboring chains, connecting the N-terminus with the MD domain of respective chains. **(e)** The DDE catalytic pocket and DNA, including the putative water molecule. **(f)** Polar interactions of R99 and R106 from the NTD1-NTD2 linker region with DNA. **(g)** Polar interactions of the NTD1 domain R77 and R81 with DNA.

### Supplementary Video Legends

**Supplementary Video 1.- Conformational variability of the shTnsB-STC complex.** The 3D variability analysis of the structure using cryoSPARC, shows that the core of the protein-DNA complex is quite stable while the arms of the X shaped DNA are highly flexible.

### Supplementary Tables

**Supplementary Table 1.**
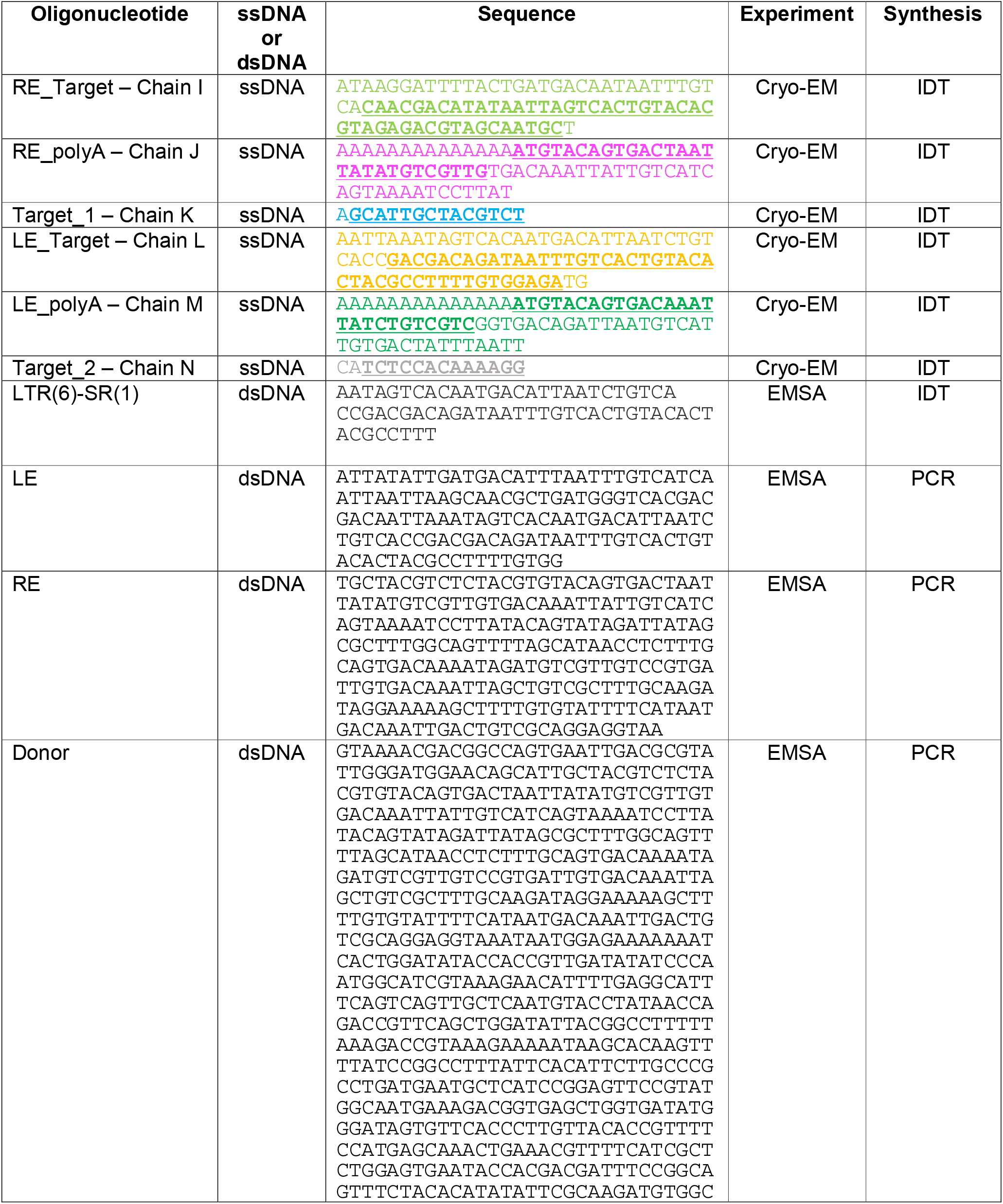
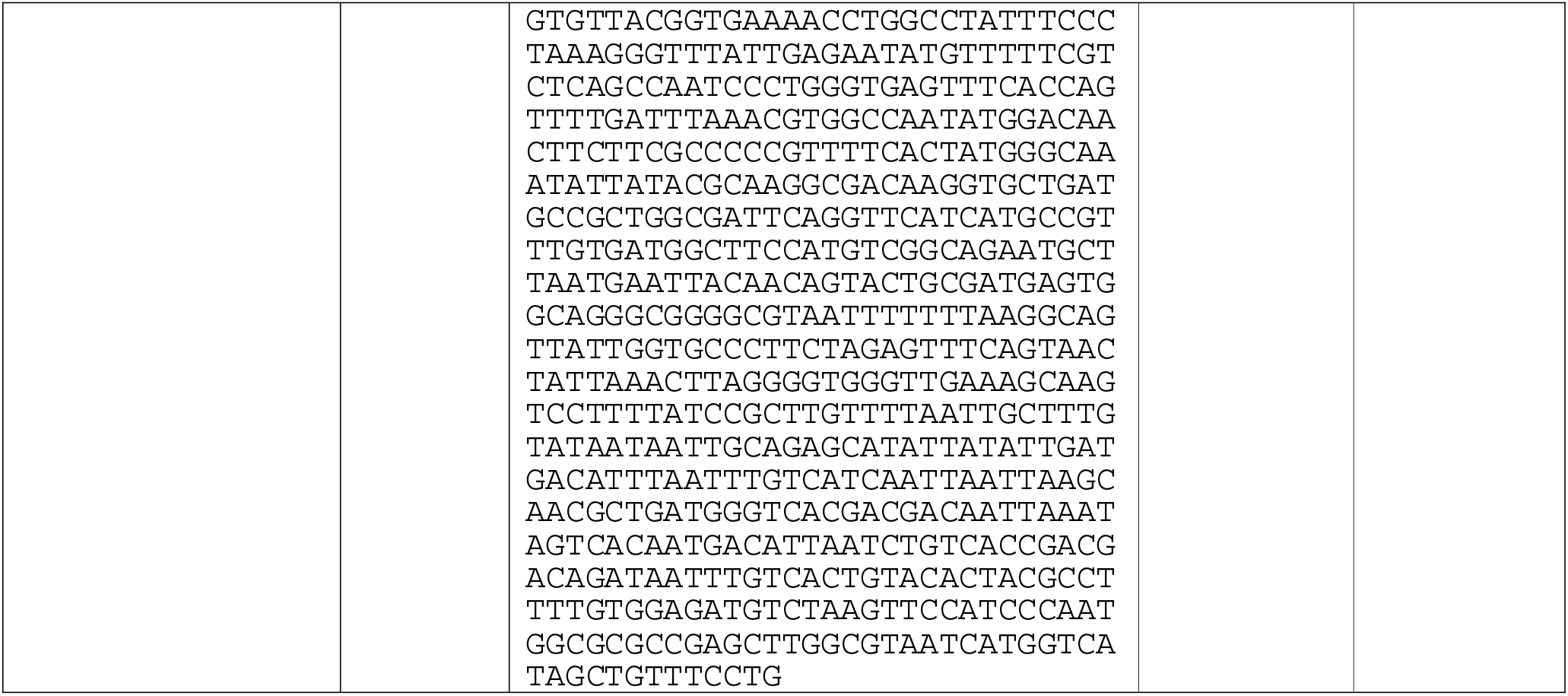
Oligonucleotides used in this study. The bold nucleotides in the DNAs used to reconstitute the shTnsB-STC complex for cryo-EM denote the sequences visualized in the structure (see also Supp. Fig. 6)

**Supplementary Table2.** Cryo-EM processing and model refinement.

|  |  |
| --- | --- |
|  | TnsB-STC |
| EMD-15294 | EMDB: |
| PDB 8AA5 | PDB: |
| **Data collection** |  |
| Electron Microscope | Titan Krios G2 |
| Voltage (kV) | 300 |
| Electron detector | Falcon III |
| Electron dose (e-/Å<sup>2</sup>) | 40 (40 frms) |
| Nominal defocus range (μm) | -0.8 to -2.0 |
| Pixel size (Å) | 0.832 |
| **3D Reconstruction** |  |
| Raw images | 4728 |
| Initial particles | 9646 K |
| Final particles | 260 K |
| Map resolution (Å), 0.143 FSC | 2.46 Å |
| Model resolution (Å), 0.5 FSC | 2.8 |
| Map sharpening B-factor (Å<sup>2</sup>) | -74.5 |
| Map sharpening method | DeepEMhancer |
| 3DFSC sphericity | 0.928 |
| Local resolution range (Å) | ~1.8 - 10 |
| **Model composition** |  |
| Protein residues | 1486 |
| Nucleotides | 172 |
| Non-hydrogen atoms | 15509 |
| Water | 2 |
| **Model refinement** |  |
| *ADP (Å<sup>2</sup>)* |  |
| Protein | 51.64 |
| Nucleotide | 47.93 |
| Water | 25.59 |
| *R.m.s. deviations* |  |
| Bond length (Å) | 0.004 |
| Bond angles (°) | 0.580 |
| *Ramachandran statistics (%)* |  |
| Favored | 95.91 |
| Allowed | 4.09 |
| Outlier | 0.00 |
| Rotamer outlier (%) | 4.62 |
| MolProbity score | 2.32 |
| Clash score | 12.42 |
| Model vs Data CC (mask) | 0.76 |

