## Supplementary figures and images for "Structure of the TnsB transposase-DNA complex of type V-K CRISPR-associated transposon"

### supp fig 1

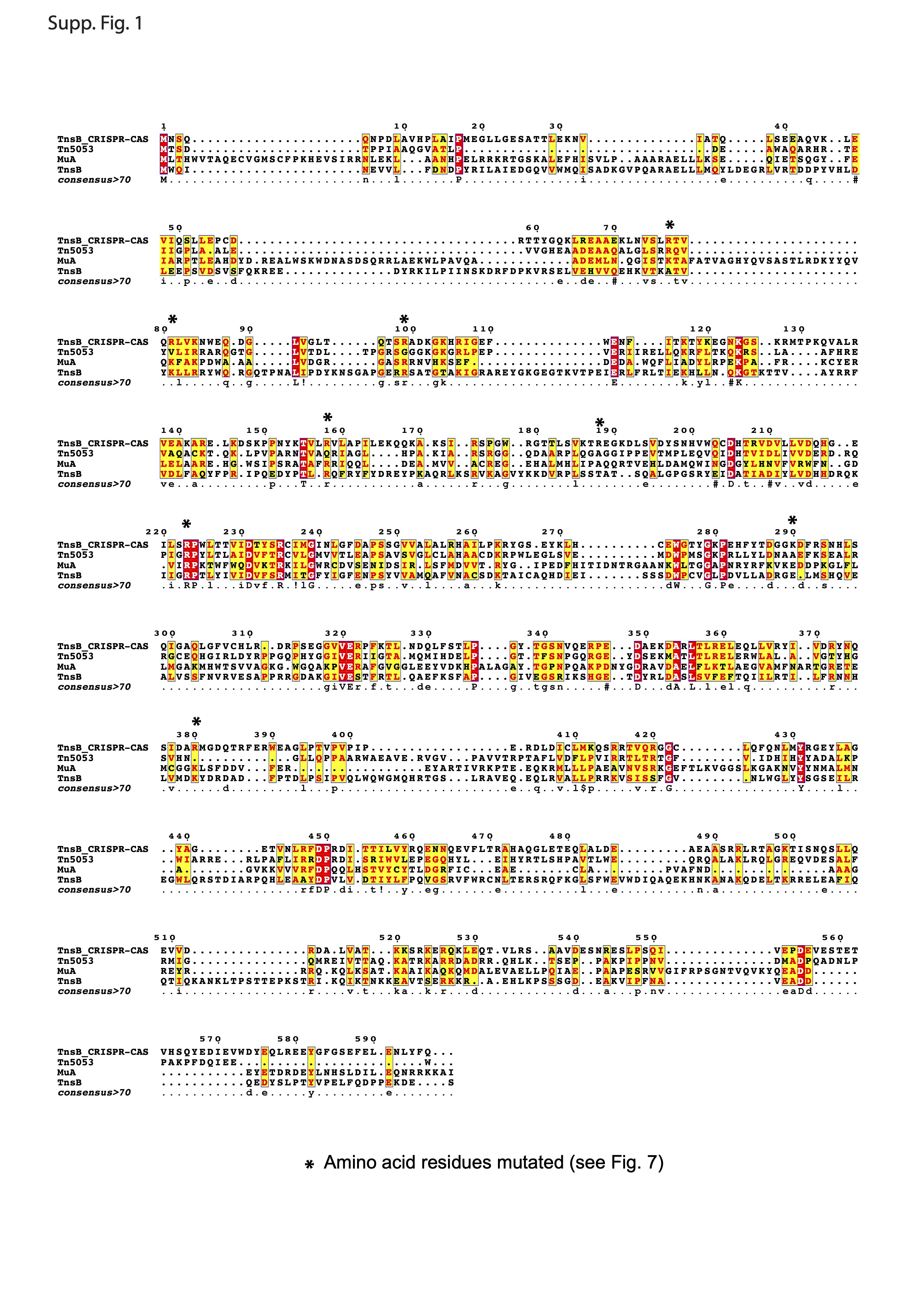

### supp fig 2

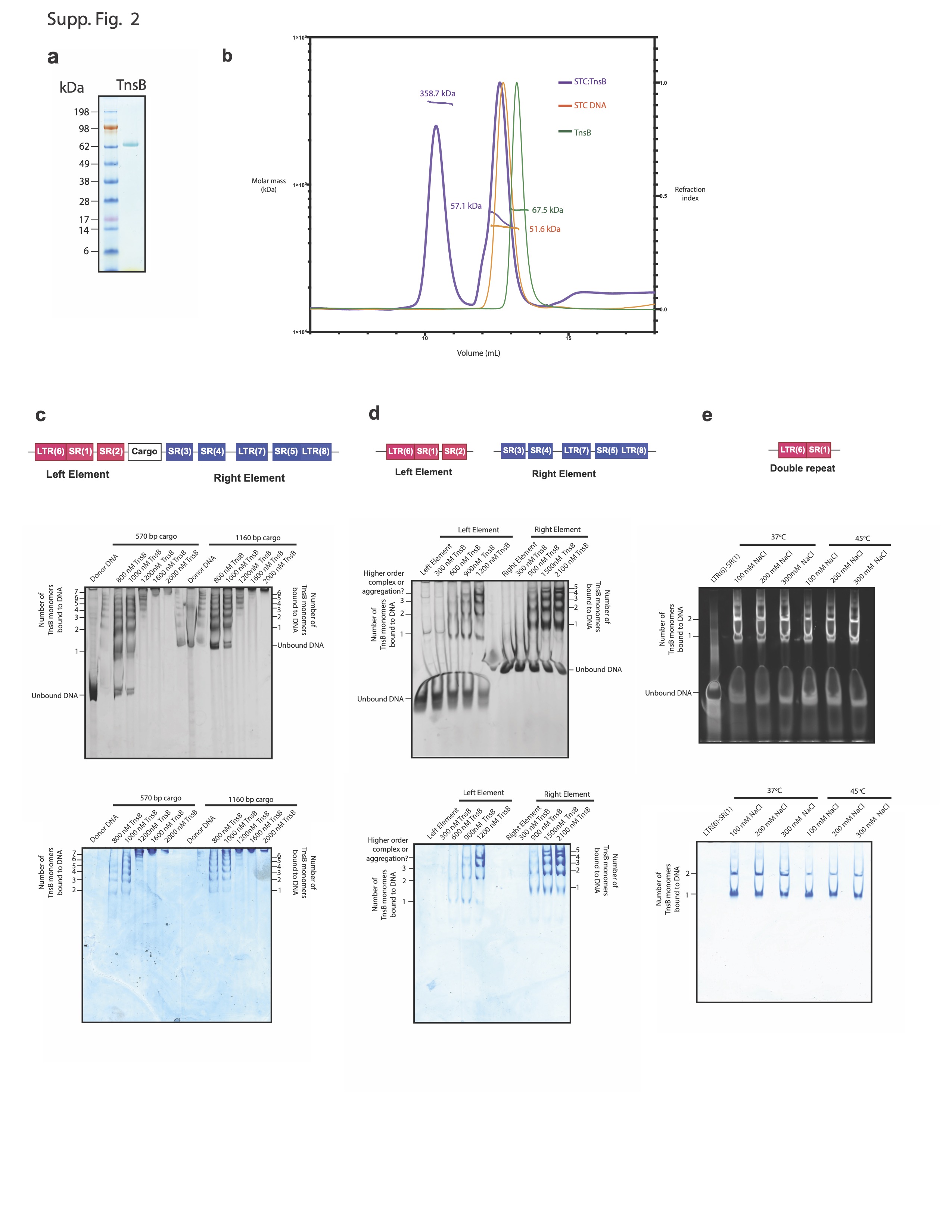

### supp fig 3

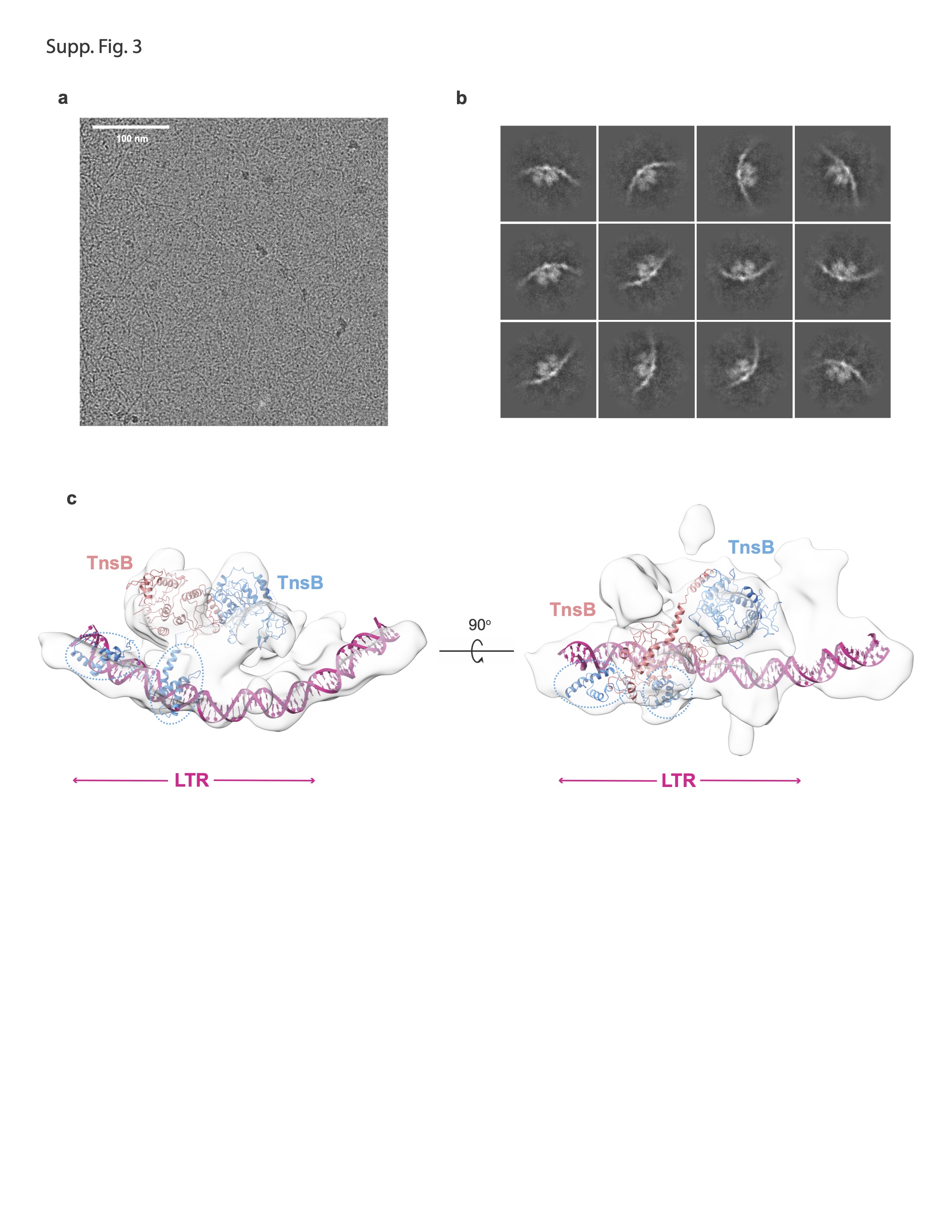

### supp fig 4

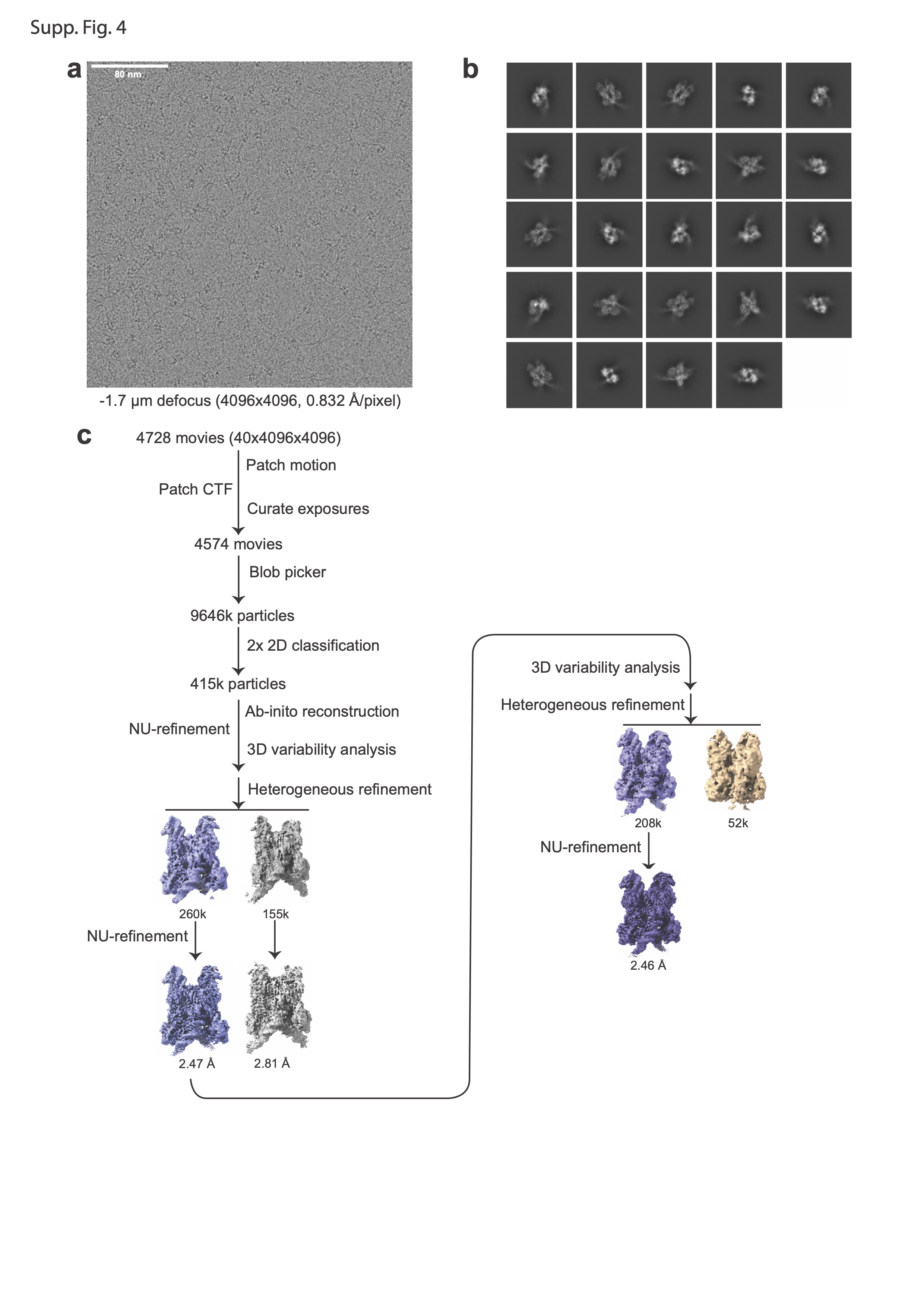

### supp fig 5

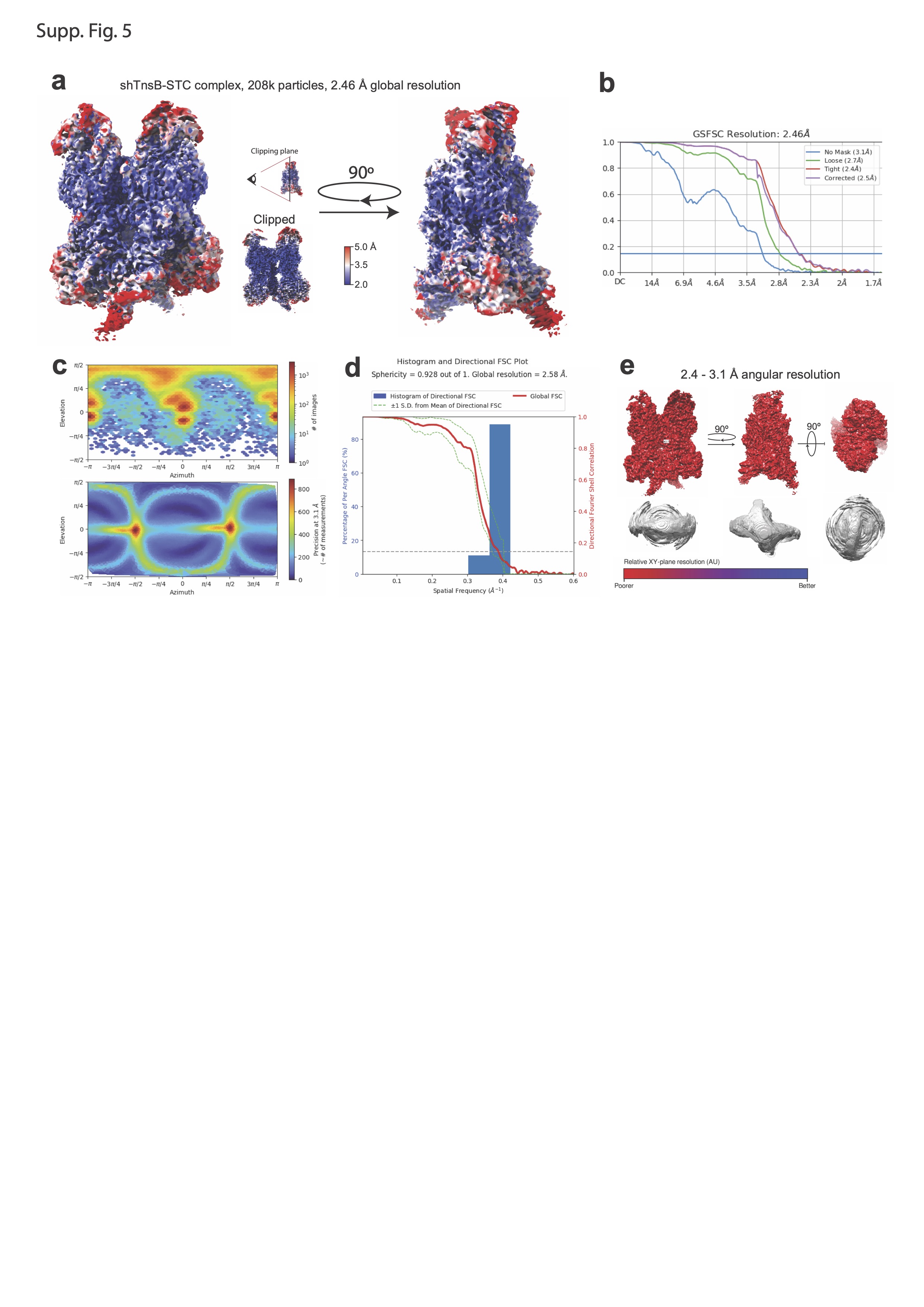

### supp fig 6

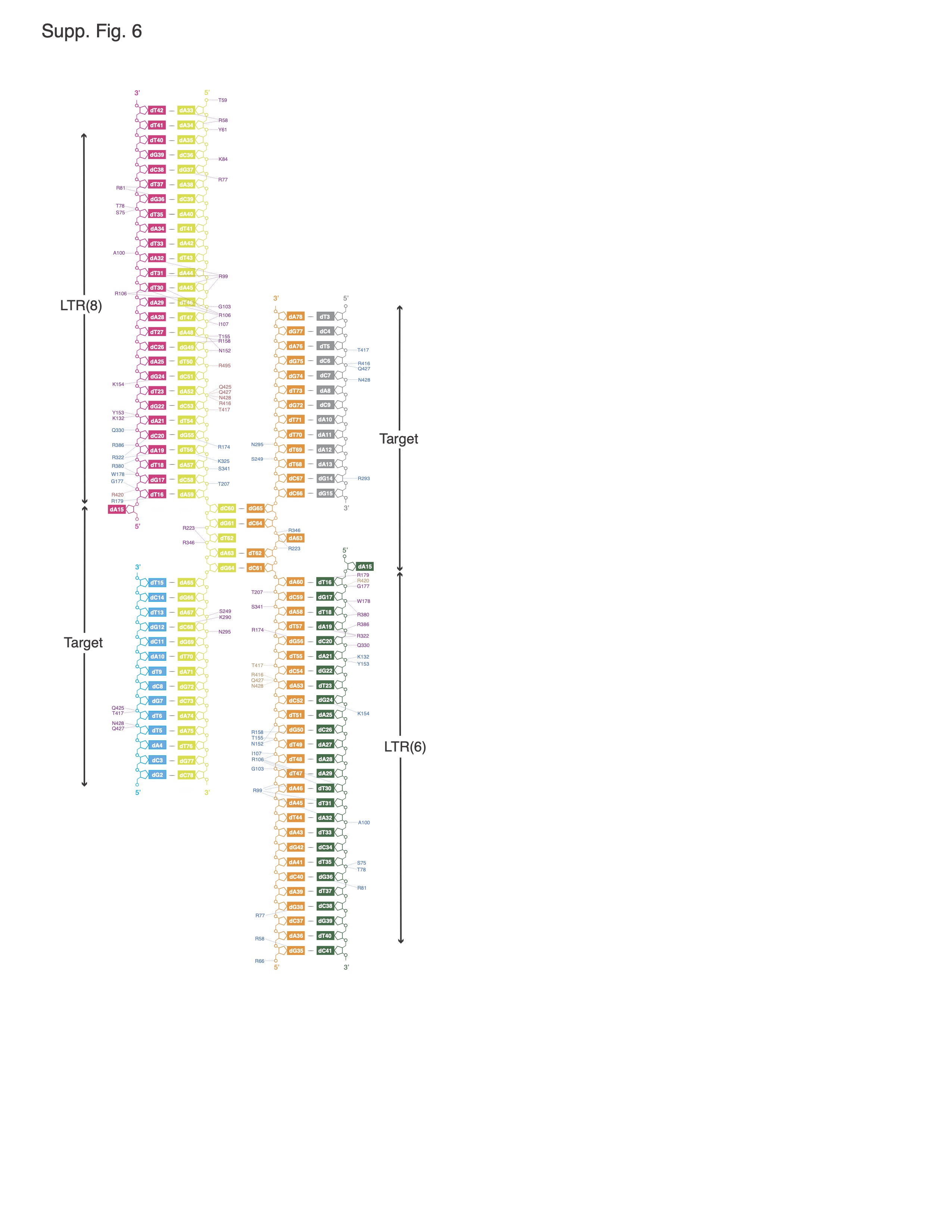

### supp fig 7

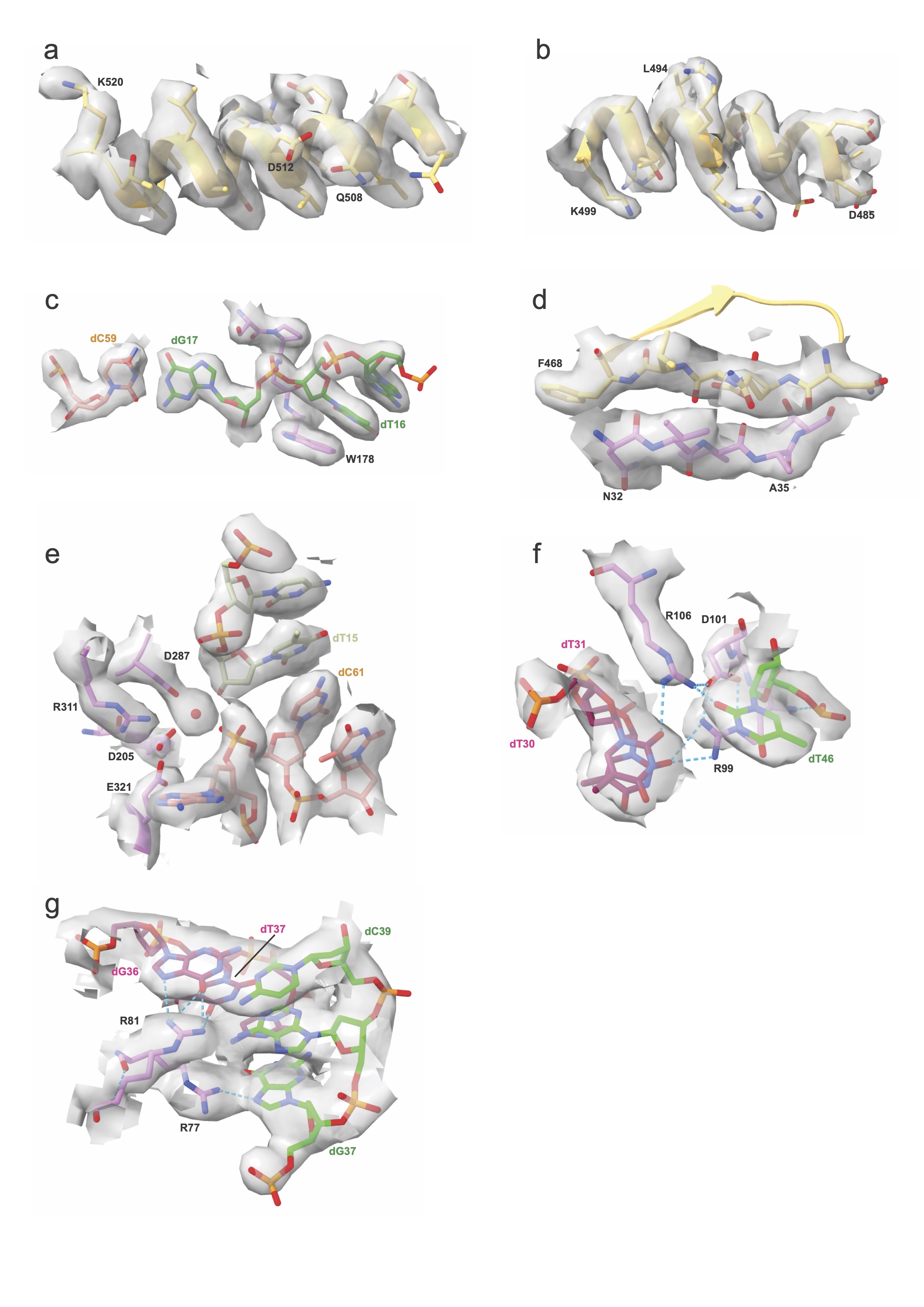
